# White-matter disconnection shapes distributed cortical spectral dynamics after stroke

**DOI:** 10.64898/2026.08.20.745699

**Authors:** Ilaria Mazzonetto, Miriam Celli, Lorenzo Pini, Antonio L. Bisogno, Giorgia Adamo, Gianluigi De Nardi, Serena De Pellegrin, Silvia Facchini, Ester Fusaro, Andrea Zangrossi, Claudio Baracchini, Anna M. Basile, Renzo Manara, Camillo Porcaro, Gustavo Deco, Maria V. Sanchez-Vives, Marcello Massimini, Maurizio Corbetta

## Abstract

Focal brain lesions are thought to induce sleep-like slow-wave activity in perilesional cortex through altered excitation-inhibition balance and structural disconnection, but whether these dynamics extend to remote yet structurally intact regions remain unclear. Here we combined source-reconstructed high-density EEG (128 channels) with structural disconnection mapping in 49 acute stroke patients and 20 age-matched controls. Cortical regions were classified as perilesional, structurally disconnected, or non-disconnected using individual lesion masks registered to normative white-matter atlases. Perilesional cortex showed increased delta and theta power and reduced beta power relative to controls. Critically, structurally disconnected regions exhibited electrophysiological changes comparable to perilesional cortex, including enhanced low-frequency activity and steeper aperiodic spectral slopes. These alterations correlated with neurological severity and multidomain behavioral impairment. Our findings demonstrate that post-stroke slow-wave activity propagates along structural disconnection pathways, providing direct electrophysiological evidence for connectional diaschisis and identifying distributed network targets for physiology-guided neuromodulation.

## INTRODUCTION

Stroke is a leading cause of long-term disability worldwide.^1,2^ The relationship between focal brain damage and behavioral impairment has been a central topic in neurology since Paul Broca’s seminal observation that speech deficits can arise from circumscribed frontal lesions. However, a purely localistic account is insufficient to explain the behavioral deficits following stroke. More than a century ago, Constantin von Monakow formalized the concept of diaschisis, proposing that focal lesions cause functional alterations in structurally intact but connected brain regions. This concept has been proved in several modern imaging studies, showing that breakdown of large-scale brain connections not directly damaged by the lesion can explain behavioral deficits and subsequent recovery.^3,4,5,6,7^ Collectively, these findings have prompted a paradigm shift in stroke, from a model of purely local damage to a network framework, characterized by changes in both local (lesional and perilesional) and distant brain (connected) regions.^8,9^ Indeed, even a relatively small lesion can alter up to 20–50% of all brain connections.^5^ Yet, the neurophysiological mechanisms underlying connectivity abnormalities remain poorly understood. In particular, the neuronal dynamics that characterize cortical regions that are structurally preserved yet disconnected following stroke are still unclear.

A long-standing clinical observation is that focal lesions produce a pronounced slowing effect on the electroencephalogram (EEG). Already in 1937, William Gray Walter reported an association between structural brain lesions and EEG slowing, most prominent in the delta frequency range, in awake subjects. Slow oscillations following stroke can persist at a chronic stage and are maximal at the location of the lesion.^10^ More recently, a mechanistic interpretation of post-stroke EEG slowing has emerged from parallels with sleep physiology.^11^ Research on non–rapid eye movement (NREM) sleep has shown that slow waves reflect profound changes in cortical excitability, characterized by bistable transitions between depolarized (ON-periods) and hyperpolarized (OFF-periods) states, and are accompanied by a breakdown of effective connectivity across cortical regions.^12^ Transcranial magnetic stimulation combined with electroencephalography (TMS-EEG) studies in sleeping and anesthetized humans have directly demonstrated that cortical bistability and the associated OFF-periods disrupt causal interactions between brain regions, providing a mechanistic link between slow oscillations and impaired information integration.^13,14^ This functional disruption is quantified by a collapse of the Perturbational Complexity Index (PCI), a measure of the spatiotemporal richness of cortical responses to direct stimulation.^15^ Critically, these effects occur in structurally intact circuits and are fully reversible upon awakening. In line with this interpretation, recent work has hypothesized that focal brain injury may give rise to similar sleep-like dynamics during wakefulness (for a review see ^16^). Importantly, connectional diaschisis (i.e. network-level abnormalities) represents a largely unexplored mechanism linking focal lesions to remote electrophysiological abnormalities.

Converging evidence from perturbational and observational approaches supports this framework. Two independent TMS-EEG studies of unilateral ischemic stroke patients,^17,18^ have revealed exaggerated slow-wave responses in perilesional cortex, indicating a local failure to sustain normal, wake-like cortical dynamics. Notably, these perilesional OFF-periods were linked to stroke severity, and their reduction over time was associated with clinical improvement and recovery of cortical interactions.^19^ Complementary evidence from intracerebral stereo-EEG recordings in patients with radiofrequency-thermocoagulation lesions has demonstrated that intracerebral activity features prominent slow waves over the perilesional area, with similar slow waves detected at cortical sites whose connectivity patterns matched the individual distribution of long-range effective connections.^20^ Furthermore, computational work has demonstrated that slow-wave dynamics are not purely the result of intrinsic cellular mechanisms but are strongly conditioned by the excitability landscape of the broader cortical network, with changes in neuronal adaptation propagating to neighboring populations through long-range pathways.^21^ In addition, preliminary TMS-EEG single-case evidence has suggested that similar abnormalities may occur outside the immediate vicinity of the lesion,^22^ but systematic neurophysiological evidence at the network level is still lacking. Critically, it remains unknown whether structural disconnection caused by stroke can induce pathological slow-wave activity in distant, structurally intact cortical regions, and whether such neurophysiological alterations contribute to the patients’ neurological deficits.

In this study, we directly address these open questions by combining high-density EEG with indirect structural disconnection mapping. Source-reconstructed EEG derived from subject-specific head models as combined with normative tractography-based disconnection maps to classify cortical parcels as perilesional, structurally disconnected, or non-disconnected. First, we test whether slow-wave activity is detectable in cortical regions that are structurally intact but disconnected from the lesion site. Second, we test whether the presence and magnitude of these slow waves are related to post-stroke severity and behavioral impairment across different functional domains. The workflow is depicted in **Fig. 1**. By bridging electrophysiology and structural connectivity within a single analytic framework, this work aims to move beyond descriptive biomarkers and provide mechanistic evidence for connectional diaschisis, with direct implications for neurorehabilitation.

**Fig 1.**
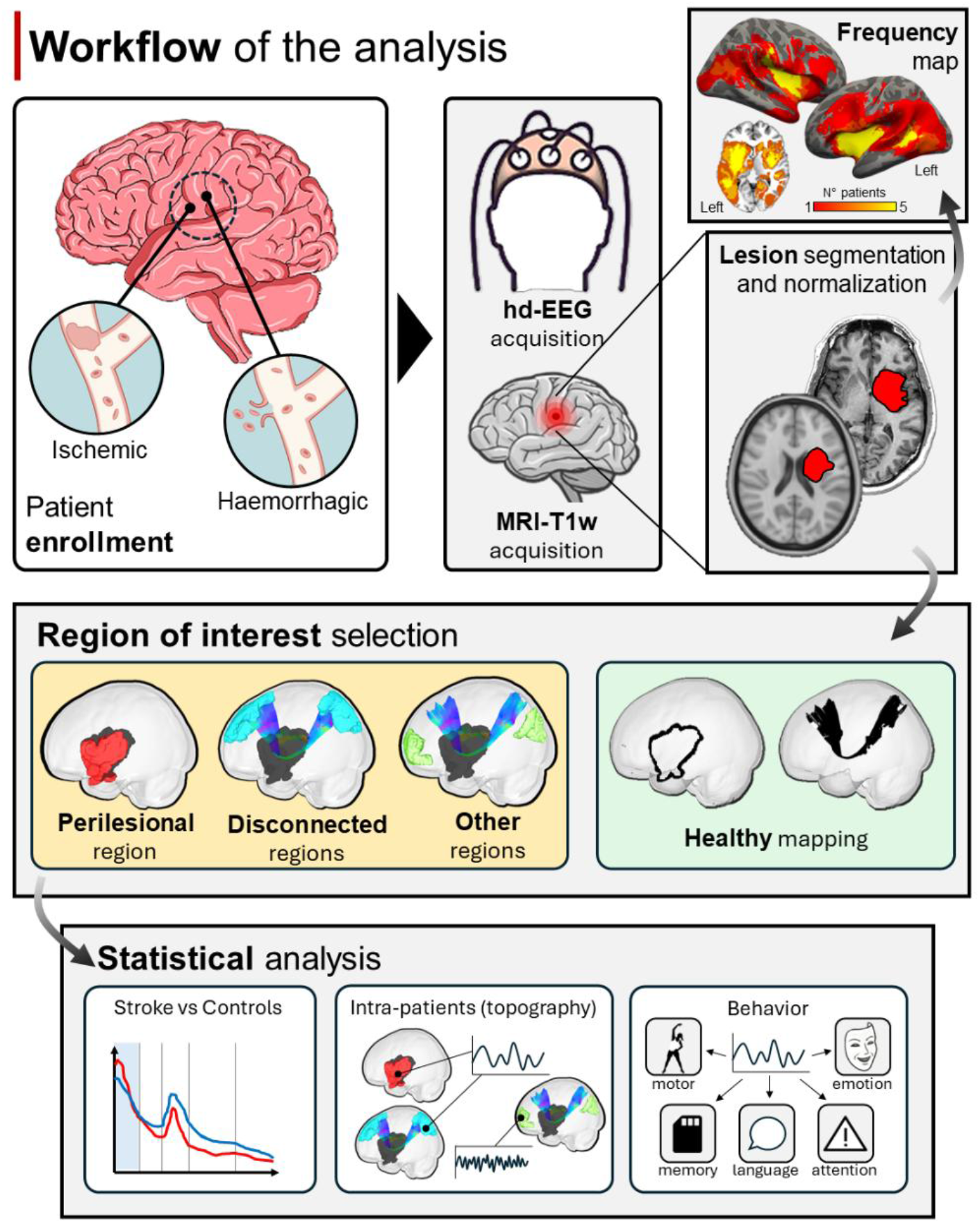
Workflow of the analysis. Acute stroke patients (ischemic or hemorrhagic) underwent high-density EEG (128-channel) and structural MRI. Lesions were segmented and normalized to standard space. Cortical regions were classified as perilesional, structurally disconnected, or non-disconnected using indirect tractography-based disconnection mapping. Age-matched healthy controls underwent the same protocol. Source-reconstructed EEG spectral power was compared between groups and across region types within patients, and associations with multi-domain behavioral outcomes were assessed. In the region-of-interest panel, an exemplary patient is shown. The perilesional region is highlighted in red (left panel), while the lesion used to compute structural disconnection is shown in black.

## RESULTS

### Participants

A total of n=88 first time stroke patients met the inclusion criteria (see methods). Of these, n=62 underwent both MRI and EEG sessions. A total of n=13 patients were excluded from the final analysis: one due to a Fazekas score of 3 on acute MRI, n=12 patients for poor EEG data quality. The enrollment process is detailed in **Fig. 2**. The selected cohort comprised n=49 stroke patients (mean age: 63±14 years; 61% male; National Institutes of Health Stroke Scale (NIHSS) score at admission: 5.8±5.8). The control group consisted of 20 healthy subjects (mean age: 65±11 years; 50% male). No significant differences were observed between stroke patients and healthy controls in age, sex, or education (p>0.46). The only differences between groups were on smokers’ proportion (p=0.04). Clinical details are reported in **Table 1**.

**Table 1.** Demographics of healthy subjects and stroke patients. P-values are shown on the right. Stroke patients showed a significantly higher proportion of smokers.

| Variable | Controls (n=20) | Stroke (n=49) | p-value |
| --- | --- | --- | --- |
| Age, mean ± SD | 65 ± 11 | 63 ± 14 | 0.87 |
| Male, % (n) | 50% (10) | 61% (30) | 0.46 |
| Education >8y, % (n) | 75% (15) | 65% (32) | 0.59 |
| Smoker, % (n) | 15% (3) | 45% (22) | 0.04* |
| Hypertension, % (n) | 35% (7) | 55% (27) | 0.32 |
| Atrial fibrillation, % (n) | 10% (2) | 12% (6) | 1.00 |
| Cardiovascular disease, % (n) | 5% (1) | 14% (7) | 0.49 |
| Diabetes, % (n) | 15% (3) | 8% (4) | 0.43 |
| Dyslipidemia, % (n) | 15% (3) | 18% (9) | 1.00 |
| **Clinical characteristics of the stroke sample** |  |  |  |
| Pre-stroke mRS, mean ± SD | | 0.12 ± 0.52 | |
| NIHSS at admission, mean ± SD | | 5.8 ± 5.8 | |
| NIHSS at discharge, mean ± SD | | 1.7 ± 2.1 | |
| mRS at discharge, mean ± SD | | 1.2 ± 1.1 | |
| Major infection |  | 1 (2) |  |
| Minor infection |  | 4 (8) |  |
| Acute heart failure |  | 1 (2) |  |
| Acute ischemic stroke (AIS) |  | 46 (94) |  |
| Intracerebral hemorrhage (ICH) |  | 3 (6) |  |
| Major infection |  | 1 (2) |  |
| Minor infection |  | 4 (8) |  |
| Acute heart failure |  | 1 (2) |  |
| Acute ischemic stroke (AIS) |  | 46 (94) |  |
| Intracerebral hemorrhage (ICH) |  | 3 (6) |  |
| **TOAST classification (AIS only, n = 46), n (%)** |  |  |  |
| Cardioembolism (CE) |  | 18 (37) |  |
| Large artery atherosclerosis (LAA) |  | 10 (18) |  |
| Small artery occlusion (SAO) |  | 2 (4) |  |
| Stroke of other cause (SOC) |  | 4 (8) |  |
| Stroke of undetermined cause (SUC) |  | 15 (31) |  |
| **ICH classification (n = 3), n (%)** |  |  |  |
| Typical |  | 2 (67) |  |
| Atypical |  | 1 (33) |  |
| **Revascularization treatment, n (%)** |  |  |  |
| Intravenous thrombolysis (rtPA) |  | 7 (14) |  |
| Endovascular thrombectomy (EVT) |  | 6 (12) |  |
| Both rtPA and EVT |  | 9 (18) |  |
| None |  | 29 (59) |  |
| **Neurological deficits at presentation, n (%)** |  |  |  |
| Motor deficit |  | 18 (37) |  |
| Language deficit |  | 13 (27) |  |
| Neglect |  | 6 (12) |  |
| **Oxford Cognitive Screen, mean (SD)** |  |  |  |
| Denomination |  | 3.48 (1.15) |  |
| Semantics |  | 2.94 (0.24) |  |
| Orientation |  | 3.94 (0.31) |  |
| Visual Field |  | 1.92 (0.34) |  |
| Sentence Reading |  | 13.4 (4.15) |  |
| Number writing |  | 2.68 (0.77) |  |
| Calculation |  | 3.56 (0.92) |  |
| Hearts Overall |  | 44.24 (8.33) |  |
| Egocentric neglect |  | -0.62 (2.55) |  |
| Allocentric neglect |  | -0.28 (0.7) |  |
| Imitating gesture dominant |  | 11.14 (2.35) |  |
| Verbal memory |  | 2.96 (1.11) |  |
| Episodic memory |  | 3.66 (0.66) |  |
| Executive function total |  | 11.42 (1.86) |  |
| Executive function mixed |  | 10.7 (4.07) |  |

**Fig 2.**
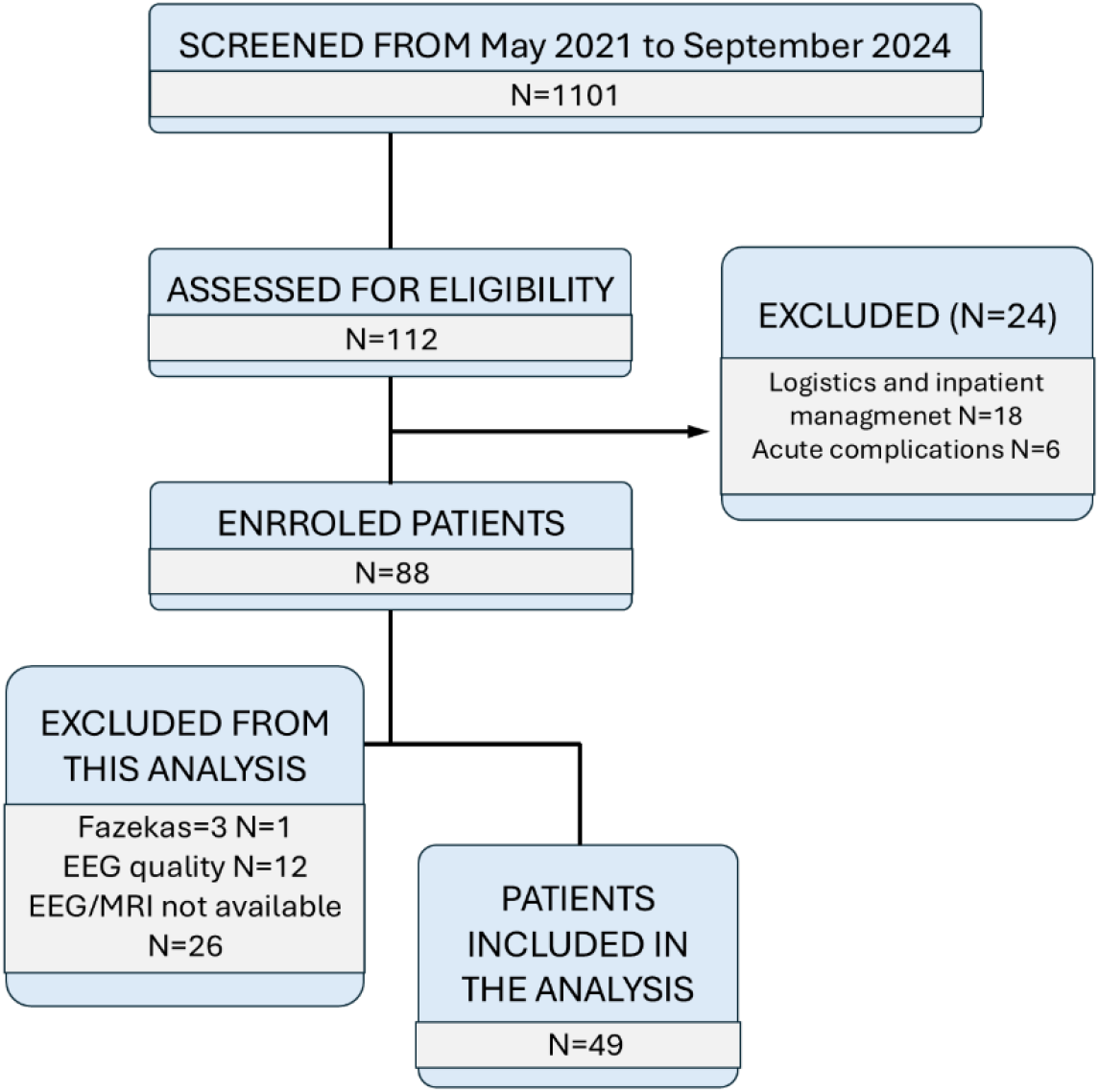
Enrollment flowchart. The figure illustrates the enrollment workflow, from patients’ screening to data used for analysis.

MRI scans were acquired on average on day 6 ± 3.5 days. EEG recordings were acquired on average on day 8 ± 2.5 days. Radiological features are reported in **Supplementary Table S1**. The distribution of lesions matched previous reports on cohorts of stroke patients recruited with similar criteria (**Fig. 1**).^34–36^

### Comparison of perilesional vs. homologous contralateral regions in stroke patients

We began by asking whether cortex immediately surrounding the lesion exhibits abnormal oscillatory dynamics relative to the intact contralateral hemisphere. First, we assessed differences between perilesional and contralateral regions in terms of power bands. A repeated measures ANOVA with a Greenhouse– Geisser correction revealed a significant effect of location (perilesional and contralateral, F(1,48)=10.36, p=0.002) and frequency band (F(2.34, 112.43)=167.90, p<0.001) and a significant location×band interaction (F(1.53, 73.84)=9.59, p<0.001), indicating that spectral profiles differed between the perilesional and contralateral regions. Post hoc comparisons showed that the perilesional region exhibited significantly higher delta power compared to the contralateral region (mean difference=0.146, p_holm_=0.001), as well as significantly higher theta (mean difference=0.118, p_ho lm_=0.001). Results are shown in **Fig. 3**.

**Fig 3.**
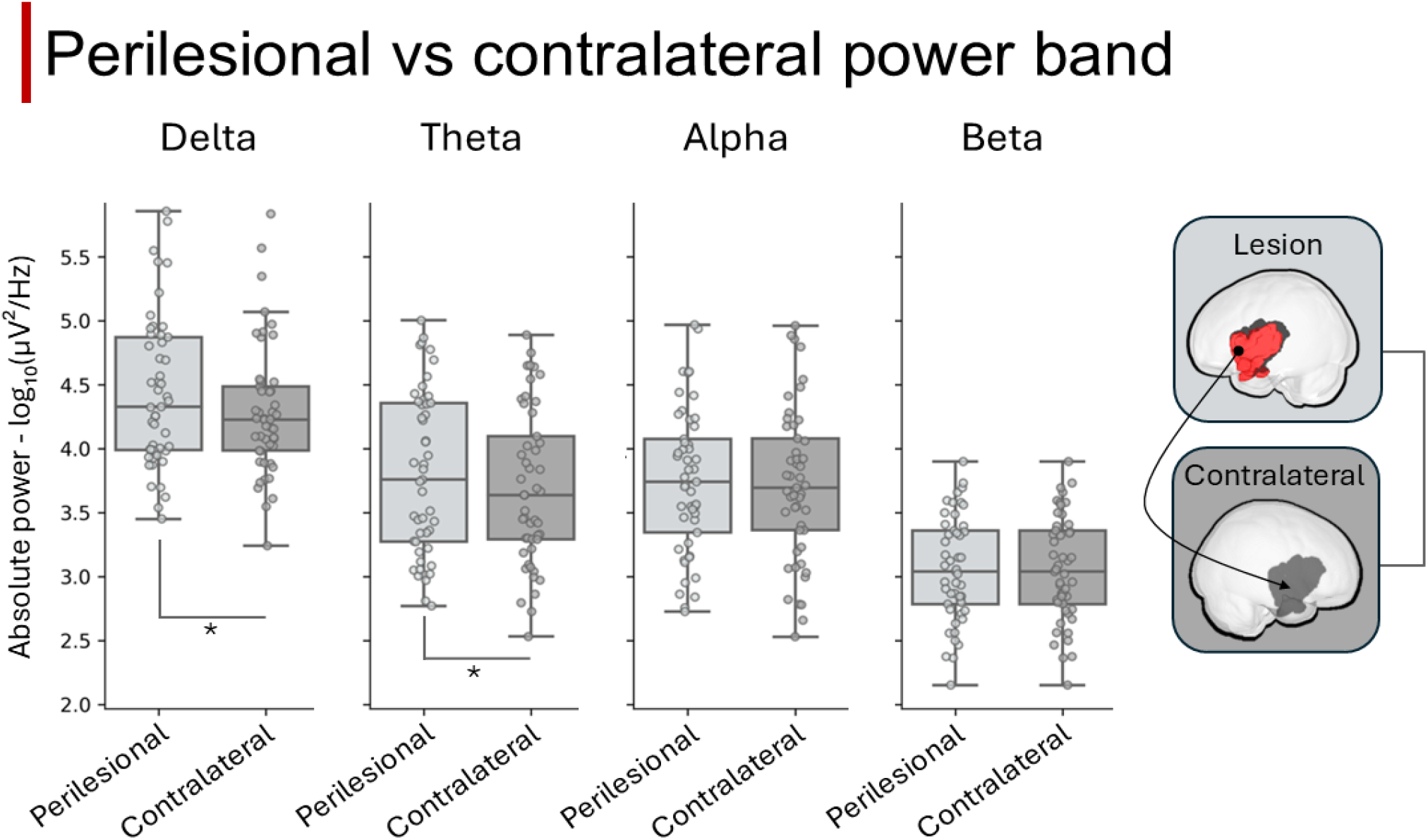
Band power differences between perilesional and contralateral regions. Differences in absolute power across frequency bands between perilesional and contralateral (flipped perilesional mask to the contralateral hemisphere) within the stroke sample. An asterisk (*) indicates the frequency bands that show a statistically significant difference.

### Comparison of perilesional regions between stroke and control subjects

We next tested whether these perilesional changes exceed normal physiological variability by benchmarking patients against healthy controls. We assessed the differences in terms of EEG band between patients and controls, within the perilesional region. A repeated measures ANOVA revealed a significant main effect of frequency band (F(1.8,175)=143, p<0.001), group (F(1,96)=55, p<0.001), and band×group interaction (F(1.8,175)=5.7, p=0.005). Post-hoc analysis in stroke patients revealed higher delta (mean difference=0.075, p_holm_=0.011), theta power (mean difference = 0.035, p_holm_=0.01) compared to controls. On the contrary, beta band was significantly lower in stroke patients compared to controls (mean difference=-0.056, p_holm_<0.001), while no significant group differences emerged in the alpha band (p=0.836). Results are shown in **Fig. 4A**.

**Fig 4.**
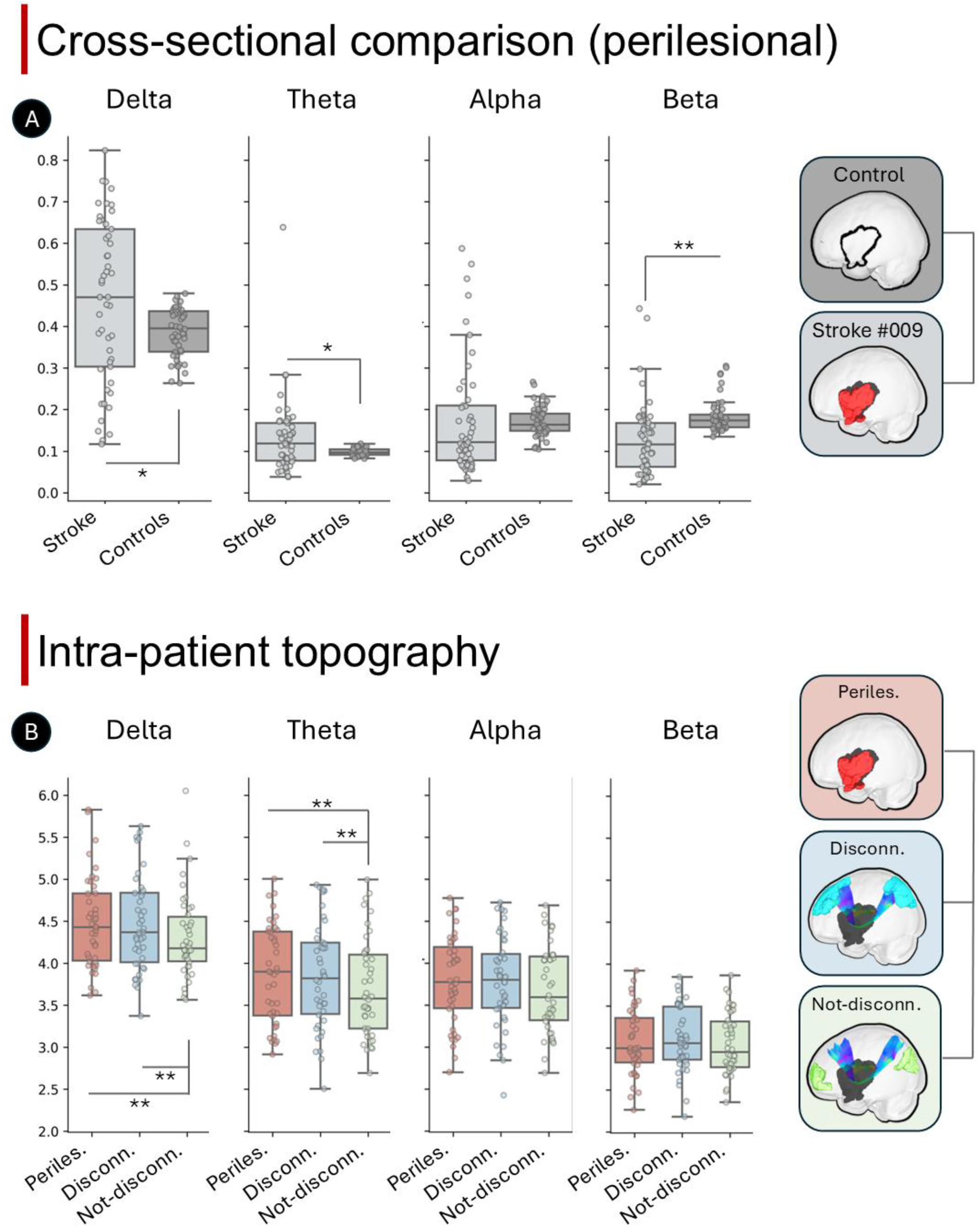
Band power differences across groups and brain regions. Panel A) Differences in relative power across frequency bands between stroke patients (red dots) and healthy controls (blue squares) in the perilesional space. An asterisk (*) indicates the frequency bands that show a statistically significant difference. Panel B) Differences in absolute power across frequency bands within the stroke patient group, across different regions: perilesional ROIs (red squares), structurally disconnected ROIs (blues squares) and not structurally disconnected ROIs (green triangles). An asterisk (*) indicates the frequency bands that show a statistically significant difference.

To capture the low-versus high-frequency imbalance in a single index, we then examined the delta/beta power ratio between patients and controls using a paired samples *t*-test. As expected, the analysis revealed a significant increase in the Delta/Beta ratio in the stroke group compared to controls (*t*(48)=3.7, *p*<0.001), suggesting a marked disruption in the balance of low- and high-frequency activity following stroke.

### Comparison among perilesional, disconnected and non-disconnected regions in stroke patients

We then addressed our central question, testing whether structurally intact cortex that is disconnected from the lesion shows the same slow-wave signature as perilesional cortex. We tested differences between perilesional, disconnected and not-disconnected brain regions within patients, to test whether regions directly disconnected by the lesion would exhibit similar perilesional activity compared to brain regions not disconnected. Results are shown in **Fig. 4B**. A repeated measures ANOVA with a Greenhouse–Geisser correction revealed a significant effect of frequency band (F(2.3, 96)=193, p<0.001), area (F(1.8,74)=7.2, p=0.002), and a significant interaction between location (perilesional, disconnected and other) and band (F(2.7, 112)=3.71, p=0.016). Post hoc comparisons showed that the perilesional and disconnected regions exhibited significantly higher delta power compared to not-disconnected regions (mean difference=0.136, p_holm_=0.009; mean difference=0.153, p_holm_<0.001, respectively). Similarly, theta in not-disconnected regions showed lower power (mean difference=0.163, p_holm_=0.001, mean difference=0.141, p_holm_<0.001, respectively). No differences emerged between regions for both alpha and beta power (p>0.093).

To complement the band-specific analysis with an aperiodic descriptor of spectral shape, we examined the spectral exponent (power slope) across the same regions. Repeated measures ANOVA revealed a significant main effect of location on power slope (F(2,82)=8.6, *p*<0.001). Post hoc pairwise comparisons (Holm-corrected) indicated that the power slope was significantly steeper (i.e., more negative) in perilesional and disconnected region-of-interests (ROIs) compared to other ROIs (*p*=0.002, and p=0.039, respectively).

### Relationship between EEG power in perilesional, disconnected and non-disconnected regions and behavioral measures in stroke patients

Having localized abnormal spectral activity to perilesional and disconnected cortex, we next asked whether these changes carry clinical significance. To this aim, participants were stratified into low- and high-power groups for each frequency band (delta, theta, alpha, and beta). For each band, 24h-NIHSS scores were compared between participants exhibiting low versus high spectral power. **Fig. 5** depicts, for each frequency band, the distribution of NIHSS scores in the two groups. In the perilesional region, the rank sum test showed that patients with higher delta power had significantly higher NIHSS scores compared to those with lower delta power (p=0.009). Lower alpha and beta power in perilesional regions was associated with higher NIHSS scores (p=0.02 and p=0.02, respectively). In disconnected regions, patients with higher delta power and lower beta power showed significantly higher NIHSS scores (p=0.03, and p=0.04, respectively). Finally, for non-disconnected regions, only beta power showed significant differences, with lower beta power associated with higher NIHSS scores (p=0.003). Results in the continuum space are reported in **Supplementary Figure S1**.

**Fig 5.**
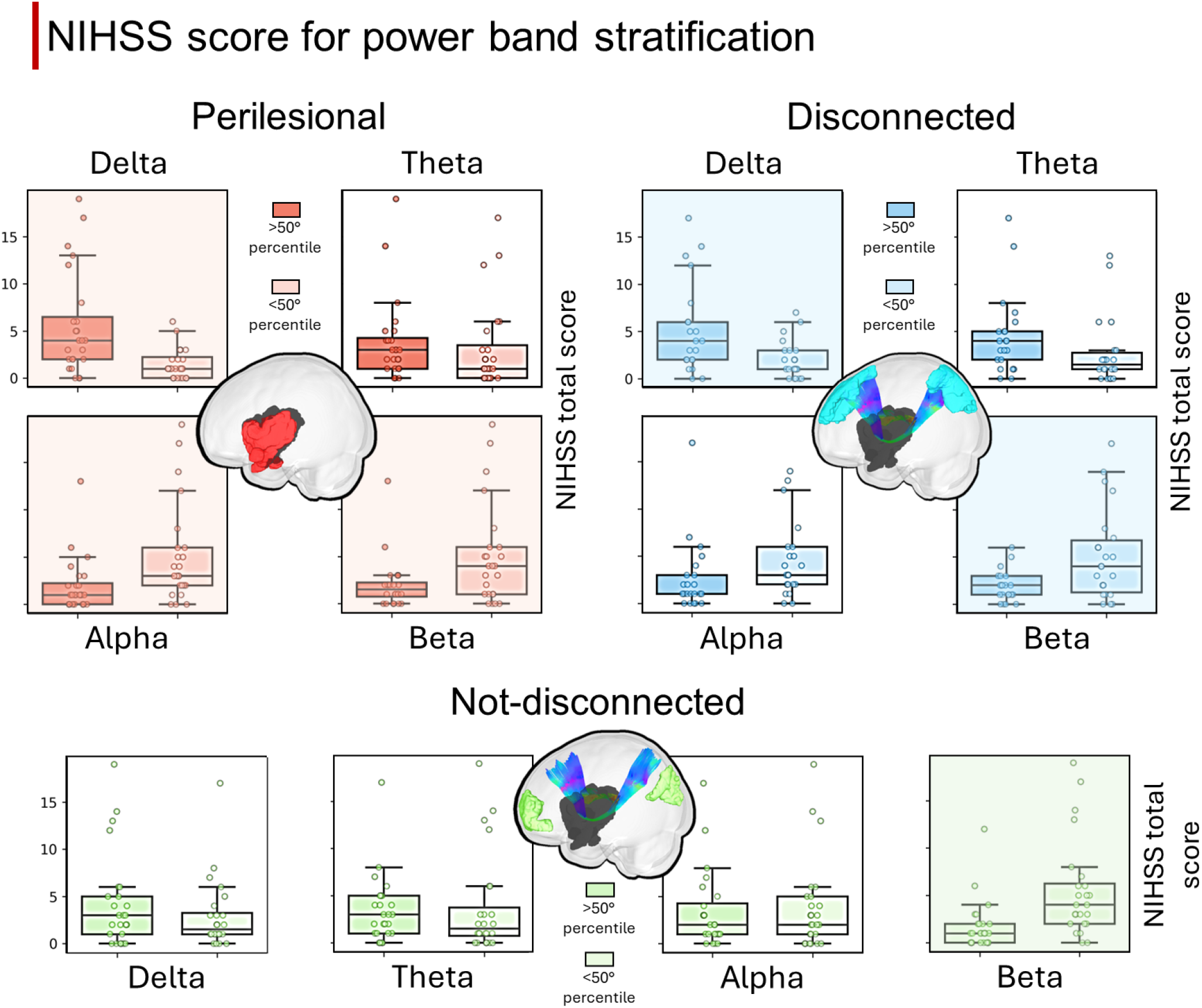
Stroke severity and power band across brain regions. Distribution of NIHSS scores in the low- and high-power groups, with power averaged across perilesional regions (top-left), disconnected regions (top-right) and not-disconnected regions (bottom). c). Statistically significant difference after FDR correction (p<0.05) are highlighted in shaded boxplots.

To assess whether these effects persisted after accounting for additional predictors, a linear regression model was fitted for each of the significant frequencies with 24h-NIHSS as the outcome and group (high/low power), age, lesion volume and acute treatment as predictors. In the adjusted models, the effect of group remains significant after adding the control variables (**Supplementary Tables S2-S7**). These findings suggest that power differences in perilesional and disconnected regions account for variance of neurological impairment (NIHSS) irrespective of many clinical variables commonly used to stratify impairment

### Relationship with behavior

To move beyond overall severity and probe domain-specific behavior, we related regional spectral power to motor and cognitive performance using a multivariate analysis. We performed a partial least square (PLS) correlation between EEG power bands (in perilesional and disconnected regions) and the four behavioral domains. The first mode (significant compared to a null model; r=0.676, p=0.002) revealed that higher cognitive performance (mainly loading on language, memory, and motor), were related to a divergent pattern across power bands, with lower delta and theta and higher alpha and beta (**Figure 6**). These results were largely confirmed by univariate Spearman’s correlation (see **Supplementary Fig S2-S5**).

**Fig 6.**
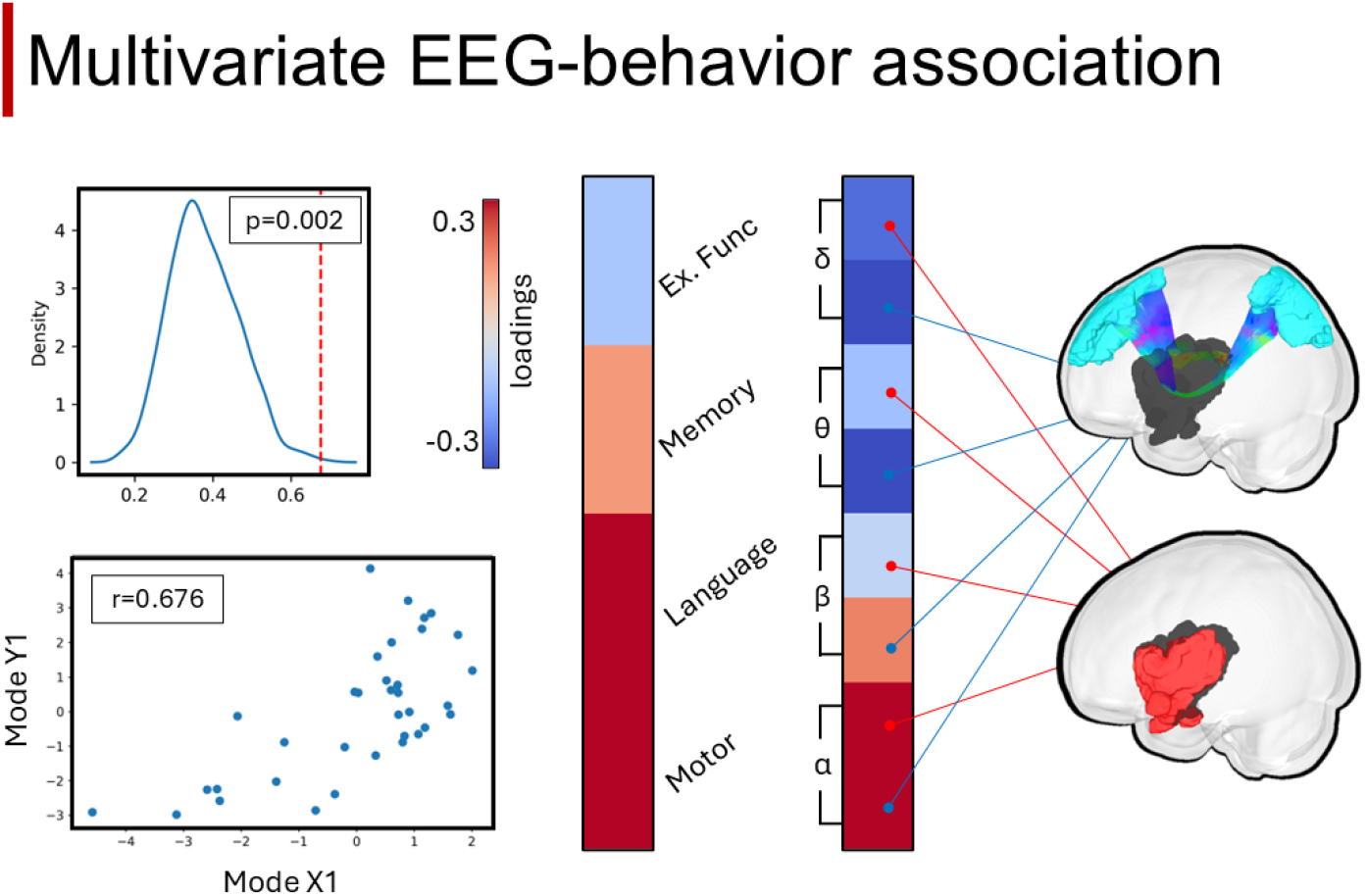
Multivariate EEG-behavioral analysis. Partial least square correlation between cognitive domains and power bands across perilesional and disconnected regions. Top left panel: comparison of the first mode r value with a null distribution; bottom-left panel: correlation between the first X and Y mode values; right panel: loadings of the first mode between cognitive and EEG power band. The arrow indicates the origin of the power band, either perilesional (red) or from disconnected regions (blue). Colors (from blue to red) indicate the loading values for PLS modes.

### Results from control analysis

We finally ran a series of control analyses to confirm the robustness of these findings, beginning with the influence of the recording condition. A first ANOVA assessed differences across bands comparing eyes-open vs eyes-closed conditions. No significant differences emerged in the delta and beta band (p>0.6), while, as expected alpha was different (p <0.001), along with a theta effect (p=0.006) (**Supplementary Table S8**).

We next verified that the results were independent of the probability threshold used to define disconnected regions. Different disconnections thresholds (40% and 60%) showed the same patterns of the main analysis (50%). Results are reported in **Supplementary Table S9-S10**.

Finally, we assessed the influence of shell size for the perilesional ROI, separately for each band. Results showed that shell size has no effect of the results reported in the main analysis (**Supplementary Table S11-S14**).

## DISCUSSION

In this study, we combined source-reconstructed high-density EEG with structural disconnection mapping to investigate how frequency-specific cortical dynamics relate to focal damage, cortico-cortical disconnection, and behavioral impairment after stroke. Our central finding is that low-frequency (delta and theta) power is increased not only in perilesional cortex but also in structurally intact regions that are disconnected from the lesion site, compared to both healthy controls and non-disconnected brain areas within the same patients. Critically, this increase in delta power was associated with greater neurological severity and poorer cognitive performance across multiple domains. These results provide direct support that structural disconnection can engage sleep-like cortical dynamics in distant, intact brain regions during wakefulness.^16^

### Mechanisms underlying slow-wave activity in disconnected cortex

The observation of increased delta activity in perilesional cortex is in line with previous literature in animal models,^37^ and with recent human TMS-EEG studies demonstrating the presence of sleep-like cortical bistability in the vicinity of ischemic lesions.^17,18^ Our results show for the first time that pathological slow-wave activity is not confined to perilesional cortex but extends to brain regions that are structurally intact yet directly disconnected from the lesion, in line with computational modeling predictions.^38^ This observation provides direct evidence that post-stroke EEG slowing follows patterns of structural disconnection at the network level.

Several neural mechanisms may underline the emergence of slow-wave activity in disconnected regions. A first possibility is that disconnection operates through a deafferentation mechanism: the loss of afferent excitatory input from lesioned or perilesional areas shifts the excitation/inhibition (E/I) balance in distant cortical targets, favoring the expression of bistable dynamics—that is, the alternation between depolarized (ON) and hyperpolarized (OFF) states that characterizes the default operating mode of deafferented cortical circuits.^39^ In this scenario, disconnected regions would display increased delta power as a proximal electrophysiological correlate of this shift in E/I balance. This interpretation is supported by Tscherpel and colleagues, who showed that damage to white matter pathways, including pedunculopontine and thalamocortical projections, is associated with TMS-evoked slow-wave activity and poorer motor outcomes after stroke,^40^ as well as by our previous observation of longitudinal structural degeneration in the main white matter tracts of stroke patients.^36^

A second, non-mutually exclusive possibility is that pathologically slow waves generated in perilesional cortex actively propagate to distant connected sites. During physiological sleep, slow oscillations behave as traveling waves that propagate along cortico-cortical and cortico-thalamocortical connections.^12,41^ Recent intracranial EEG recordings in awake brain-injured patients have confirmed that sleep-like slow waves can travel along long-range cortico-cortical pathways during wakefulness,^20^ and animal models have demonstrated a causal link between chemogenetic inactivation of a cortical area, the generation of local slow waves, and their propagation to distant connected regions with consequent large-scale functional connectivity alterations.^42^

These two mechanisms (distant deafferentation and active traveling) may coexist and reinforce each other in extending cortical bistability beyond the perilesional area. Future studies combining source-reconstructed high-density EEG with directed connectivity analyses, or employing causal perturbational approaches such as TMS-EEG, will be essential to disentangle these mechanisms.

### Electrophysiological correlates of disconnection and behavioral relevance

A core finding of this study is the association between increased delta power in perilesional and disconnected regions and clinical severity, as indexed by the 24-hour NIHSS. This time point captures the integrated effect of initial neurological impairment, acute revascularization therapies, early infarct evolution, and medical complications, thus providing a clinically meaningful summary of post-stroke status.^24^ At this time point, the NIHSS reflects the combined effects of baseline neurological impairment, acute revascularization therapies (including intravenous thrombolysis and endovascular thrombectomy), infarct evolution, and early medical complications, providing an integrated summary of clinical impact.^43^

The association between NIHSS scores and frequency-specific alterations (increased slow-wave activity alongside reduced fast-wave activity) in structurally disconnected regions suggests that remote electrophysiological changes are not merely an epiphenomenon of focal damage. Rather, they likely reflect system-level consequences of disconnection, whereby distributed network dysfunction actively shapes clinical severity.

This interpretation is reinforced by the multivariate and univariate analyses relating EEG spectral power to more in-depth motor and cognitive measures. The PLS analysis revealed that higher delta and theta power, together with reduced alpha and beta power, were associated with poorer performance across multiple behavioral domains. Notably, the loadings were slightly stronger for disconnected than for perilesional regions (although univariate analysis showed similar magnitude), suggesting that the electrophysiological fingerprint of network-level disruption may contribute to behavioral variance beyond what is captured by perilesional changes alone. These results are consistent with previous evidence showing that altered local and global connectivity patterns, rather than lesion topography per se, explain behavioral deficits and recovery trajectories after stroke.^3,7,44–47^

Interestingly, we also found a reduction of beta activity in non-disconnected regions that may reflect altered information processing also in not-directly disconnected cortex. In fact, previous fMRI connectivity work showed alterations not only in first order (regions directly connected to/from the lesion site), but also higher order connections (regions indirectly connected through second and higher order interactions).^44^

### Clinical implications

The presence of pathological slow-wave activity in the structurally intact but disconnected cortex suggests that these regions remain viable substrates for intervention. Targeting such areas may allow restoration of more physiological cortical dynamics and enhance the effectiveness of rehabilitation strategies. Furthermore, slow waves and decrements of high frequency EEG activity may become an important proxy for very early diagnosis (e.g. on an ambulance) or for prognosis prediction.

Accumulating evidence indicates that slow waves can be influenced by pharmacological approaches and optogenetics in experimental models, in vivo and in vitro.^48,49^ In this context, emerging new non-invasive technologies, such as transcranial ultrasonic stimulation, offer high deep-brain selectivity.^50^ The design of effective stimulation protocols, however, critically depends on whether post-stroke slow waves represent a transient, homeostatic response to injury or a maladaptive process that actively constrains recovery. Evidence from healthy subjects suggests that slow-wave activity emerging in deafferented motor cortex may reflect a compensatory homeostatic mechanism that helps disconnected neurons remain active despite reduced input.^51^ Cassidy and colleagues found delta band power related to greater injury and better motor status in chronic stroke patients.^52^ Consistently, during physiological sleep, slow waves are widely regarded as protective and restorative.^52^ Future longitudinal studies will therefore be necessary to determine whether sleep-like dynamics during wakefulness serve local protective or restorative functions analogous to forms of “local sleep,” or instead interfere with network integration and impair neuroplastic recovery.

### Limitations and strengths

Several limitations should be acknowledged. First, the cross-sectional design precludes causal inferences about the temporal evolution of electrophysiological and structural changes. Longitudinal assessments will be necessary to determine whether reductions in delta activity parallel structural recovery and behavioral improvement, and whether slow-wave dynamics can serve as dynamic biomarkers of recovery. Second, although our findings are consistent with the cortical bistability framework,^16^ resting-state EEG alone cannot directly demonstrate the presence of bistable OFF-periods. Future work integrating diffusion imaging with causal perturbational approaches such as TMS-EEG^17,18^ will be essential to directly test the disconnection-bistability hypothesis at the individual patient level. Third, the sample size, although comparable to previous multimodal stroke studies, may limit the detection of subtler effects, particularly in subgroup analyses.

## CONCLUSIONS

In summary, by combining source-reconstructed high-density EEG with structural disconnection mapping, we provide direct electrophysiological evidence that post-stroke slow-wave activity extends beyond perilesional cortex to structurally intact brain regions following patterns of white-matter disconnection. These network-level pathological dynamics are behaviorally relevant and offer a mechanistic account of connectional diaschisis. As such, they represent candidate neurophysiological targets for prognostic biomarkers and network-informed neuromodulatory interventions aimed at promoting stroke recovery.

## METHODS

### Study design and participants

From May 2021 to September 2024, patients were prospectively recruited from the Neurology Clinic and Stroke Unit of the University of Padova Hospital, as well as the Stroke Unit of S. Antonio Hospital in Padova. Recruitment involved individuals who had experienced a first symptomatic stroke (ischemic or hemorrhagic) and met defined inclusion and exclusion criteria.

Patients were recruited between 72 hours and 2 weeks after the event if i) they were 18 years of age or older (no upper age limit), ii) had experienced a first symptomatic ischemic or hemorrhagic stroke, and iii) were awake, alert, and capable of active participation. Exclusion criteria comprised inability to maintain wakefulness during testing; stroke affecting multiple vascular territories; the presence of neurological, psychiatric, or medical comorbidities that could interfere with study participation or data interpretation (e.g., dementia, schizophrenia) or substantially limit life expectancy to less than one year (e.g., advanced cancer, New York Heart Association class IV congestive heart failure); clinically significant periventricular white matter disease (Fazekas=3); more than two lacunes, clinically silent, less than 15 mm in size on admission CT scan; claustrophobia; and confirmed or suspected mobile metallic implants or fragments (including intraocular) precluding MRI. We additionally recruited a sample of healthy participants. Healthy control participants were adults (≥18 years) in good health, comparable to the stroke cohort in age, sex, and typically recruited from patients’ relatives. Exclusion criteria for controls included prior stroke, central nervous system tumors, history of dementia, prior neurosurgery, severe psychiatric illness, or inability to provide informed consent.

All enrolled patients underwent a comprehensive assessment within two weeks of stroke onset. A second evaluation identical to the first one was carried out at three months (not reported in the present study). The evaluation included a neurological examination, a speech and neuropsychological evaluation, high-density EEG recording and MRI acquisition (described below). Healthy controls underwent the same neurological cognitive and imaging examination.

The procedures outlined in this study were approved by the Ethics Committee of the AOPD (Study AOP1995, code CESC 4996/AO/21, approval protocol 22903/2021).

### Behavioral assessment

Neurological severity was quantified using the National Institutes of Health Stroke Scale (NIHSS)^23^. The NIHSS was administered upon admission, at 24 hours and at approximately 1 week after stroke onset. The 24-hour NIHSS score (24h-NIHSS) was used for subsequent analyses, as it represents one of the most reliable and consistent predictors of early stroke severity and functional outcome.^24^

Behavioral assessment was conducted at the time of EEG acquisition and consisted of a comprehensive battery of standardized neuropsychological tests together with the Oxford Cognitive Screen.^25^ The assessment required approximately 90 minutes. The following instruments were analyzed in this study: the Frontal Assessment Battery (FAB),^26^ the Rey Auditory Verbal Learning Test (RAVLT),^27^, the Brief Visuospatial Memory Test (BVMT),^28^ the Trail Making Test (TMT, version A and B),^29^ and the Aphasia Test (ENPA).^30^ See Supplementary Methods and Supplementary Figure S6 for a complete description.

### MRI Acquisition

Imaging data were acquired using a 3T Philips Ingenia whole body MR scanner equipped with a 32-channel headcoil, at the Neuroradiology Unit of the University Hospital of Padova, Italy. High-resolution anatomical images were obtained using the following sequences: (1) 3D T1-weighted Magnetization Prepared Rapid Acquisition Gradient Echo (MPRAGE) sequence (echo time/repetition time/inversion time = 2.36/1700/1000 ms, 1 mm slices, 240 mm field of view, 256 × 256 acquisition matrix); (2) 3D Fluid-attenuated inversion recovery (FLAIR) (repetition time/echo time/inversion time = 5000/394/1800 ms, 1 mm^3^ isotropic resolution); and (3) 3D T2-weighted sequence (repetition time/echo time = 3200/564 ms, 1 mm^3^ isotropic resolution). Multi-shell diffusion images and resting-state functional MRI were also collected but not included in the present study.

### Region of interests

Subacute lesions following stroke were segmented from FLAIR and DWI scans. All lesions were manually segmented by a board-certified neurologist (ALB) using ITK-SNAP tool software.^31^ Fazekas scores and number of lacunes in CT admission scans were also quantified. Spatial transformations were estimated using ANTs. Specifically, we computed an affine transformation combined with a bi-directional diffeomorphic registration to align individual brains to MNI space. This procedure was applied to both stroke patients and healthy controls. By combining the transformation from patient space to MNI space with the inverse transformation from control space to MNI, each lesion mask was mapped onto the brains of all control subjects.

Each patient’s lesion mask was registered to the MNI template using the spatial transformation previously estimated with ANTs. This procedure provides, for each voxel, the probability of being traversed by a fiber that would be disconnected because of the lesion. Disconnection probabilities were computed for each gray matter parcel based on the Brainnetome Atlas and the SUIT atlas, resulting in a total of 274 ROIs.

We used the BCB toolkit^32^ to estimate indirectly the structural disconnection caused by a lesion. The inference about the affected structural pathways is made by embedding the lesion into a normative structural connectome obtained from a sample of healthy subjects. The lesions were normalized to MNI space resampled to 1 × 1 × 1 mm, to match the space of the tracts included in the BCB toolkit. For each voxel, the likelihood that a white matter bundle directly connected with the lesion passes through it is calculated.

We defined four ROI categories (three parcel-based or comparison within patients, and one voxel-based for cross-sectional analysis, i.e., patients vs controls): i) perilesional: parcels intersecting a region extending up to 2.4 cm from the lesion border (from 4 to 20 mm); ii) disconnected: parcels with a probability of disconnection greater than 50%, excluding those classified as perilesional; iii) not-disconnected: parcels with a disconnection probability below 10% and not included in the perilesional space; iv) perilesional voxel-based region defined as the surrounding voxels ranging from 4 to 20 mm from the lesion border and the corresponding contralateral regions created flipping the perilesional voxel space. Voxels from the contralateral region encompassing the perilesional masks (and viceversa) were eliminated to avoid overlapping voxels between the two masks.

### EEG acquisition and preprocessing

We recorded two sessions of high-density EEG, consisting of 10 minutes with eyes open and 10 minutes with eyes closed. The session order was randomized across participants. Individual high-resolution T1-weighted structural MRI scans were imported into the Nexstim Ltd. Navigation System and initially co-registered with digitized anatomical landmarks. Whole-scalp EEG data were acquired using a 128-channel Bittium amplifier system (Brain Products GmbH), with electrodes positioned according to the extended 10–20 International System. Electrode impedances were maintained below 10 kΩ, and signals were sampled at 1,000 Hz. Digitized electrode locations were co-registered to individual MRI scans using the navigation system.

EEG data were pre-processed using custom Matlab (The MathWorks, Inc, Natick, Massachusetts, USA) scripts based on functions from the EEGLAB software (version 2022.0, 70). First, data were filtered with a set of three different FIR filters: 1) a low-pass filter with a cut-off frequency of 1 Hz; 2) a high-pass filter with a cut-off frequency of 80 Hz; 3) a notch filter with cut-off frequency at 49-51 Hz. The FIR filter was implemented with eeglab pop_eegfiltnew function with default settings. Afterwards EEG data were resampled at 250 Hz. Then, an automated detection of the noisy channels was performed and later confirmed by visual inspection. Selection was based on the combination of the following five criteria, whose thresholds were determined with a preliminary examination of the dataset to optimize the detection: i) impedance at the end of the acquisition above 20 kΩ; ii) absolute amplitude bigger than 650; iii) standard deviation bigger than 7 for the improbability test, iii-iv) standard deviation bigger than 4 for the spectral test; v) standard deviation bigger than 4 for the kurtosis test. Channels selected by the criterium i), ii) or at least two of iii-v) criteria were interpolated using spherical splines. Selected channels were also visually checked to ensure accuracy of the automated detection. Data were then re-referenced to the average of all channels. Ocular, muscular, movement and noise artifacts were removed by applying Independent Component Analysis (ICA) using a deflation-based Fast fixed-point ICA algorithm with the hyperbolic tangent as cost function. Artifactual independent components (i.e., muscle, eye, heart, channel noise and line noise components) were manually selected after visual inspection. Components computed on the 1-80 Hz filtered dataset were then removed from a dataset filtered with a set of three different FIR filters: 1) a low-pass filter with a cut-off frequency of 0.2 Hz; 2) a high-pass filter with a cut-off frequency of 80 Hz; 3) a notch filter with cut-off frequency at 49-51 Hz. In this final dataset the original sampling frequency of 1000 Hz was kept and all the bad channels identified in the previous steps were removed before component rejection.

### EEG Source Reconstruction

The EEG data were projected into source space to estimate the underlying neural generators. To address the source localization problem, we first solved the forward problem, which involves calculating the scalp potentials generated by current dipoles located within the brain. Subsequently, the inverse problem was tackled by estimating the neural sources that best explain the observed scalp potentials, based on the recorded EEG data.

A subject-specific head model was constructed by segmenting the T1-weighted image of each participant into five-tissue types: white matter, gray matter, cerebrospinal fluid, scalp, and bone. Segmentation was performed using SPM12 after applying bias-field correction using the N4 algorithm implemented in ANTs. Tissue conductivity values were assigned based on established literature.

EEG electrode positions, obtained via digitization, were already aligned to the T1-weighted image. To ensure accuracy and avoid digitization errors (e.g., duplicated or missing electrodes, or incorrect label assignments), we first computed a rigid transformation based on anatomical landmarks (nasion, inion, and left/right preauricular points) between the template cap and the subject’s space. We then applied an affine Coherent Point Drift registration between the digitized electrode positions and the template configuration. Electrode pairs with Euclidean distances exceeding the average were corrected by replacing the digitized position with the corresponding location from the registered template. The final electrode positions were then projected onto the scalp surface mesh to ensure accurate anatomical alignment.

Dipole source locations were defined by discretizing the gray matter into a regular 3D grid with 4 mm spacing. Dipoles located within damaged areas were excluded. Lesion masks, defined in FLAIR space, were coregistered to the T1-weighted MRI using a rigid transformation, then resampled to match the resolution of the source space.

For some participants, sensor digitization data was not available. In these cases, a cap template was employed. The template was first rigidly aligned to the scalp mesh using anatomical landmarks, and then further refined using the Coherent Point Drift algorithm. When the T1-weighted image was unavailable, stroke patients were excluded from the analysis, as it would not have been possible to construct a subject-specific head model and to exclude dipoles located within the lesioned region. In contrast, for control participants, the MNI template was used without such limitations.

At each grid point, three orthogonal unit dipoles (x, y, z directions) were modeled. The forward solution, which describes how neural sources generate scalp EEG signals given to the head model, was computed using the Simbio finite element method as implemented in the FieldTrip toolbox. The inverse problem was solved using the exact Low Resolution Brain Electromagnetic Tomography (eLORETA) algorithm, also implemented in FieldTrip. For each source location, the first principal component across the x, y, and z directions was extracted to represent the estimated neural time series.

Power spectral densities (PSD) of each source time series were computed using Welch’s method, with 2-second epochs and 50% overlap. The resulting spectra were segmented into four standard frequency bands: Delta (0.5–4 Hz), Theta (4.5–8 Hz), Alpha (8.5–13 Hz), and Beta (13.5–30 Hz). For each band, both absolute and relative power were calculated. The Delta/Alpha and Delta/Beta ratio were also computed.

Following the approach described by Donoghue,^33^ we also extracted the slope of the aperiodic (1/f-like) component of the power spectrum, as it may provide a more concise summary of spectral changes, particularly in cases where low-frequency power increases and high-frequency power decreases as a consequence of the lesion.

### Comparison of perilesional vs. homologous contralateral regions in stroke patients

Each lesion mask was flipped along the left-right axis. For each mask configuration, we computed the mean power across the different frequency bands. A repeated-measures ANOVA was performed to assess the within-subject effects of the factors band (delta, theta, alpha and beta) and side (perilesional vs contralateral). Greenhouse-Geisser or Huynh-Feldt corrections were applied in cases of sphericity violations (Mauchly’s test. Post hoc comparisons, corrected using Holm’s method, were conducted to identify the frequency bands in which power significantly differed between the contralateral and perilesional regions.

### Cross-sectional perilesional differences between stroke and controls

For each patient, we calculated the mean power within the perilesional space (voxel-space) across different frequency bands, as well as the mean power in the corresponding regions of the control subjects.Repeated-measures ANOVA was conducted using JASP to assess the within-subject effects of the factor band (delta, theta, alpha and beta) and the between-subject effects of group (stroke vs controls). Violations of the sphericity assumption were tested using Mauchly’s test; when violations were detected, the degrees of freedom were adjusted using either the Huynh-Feldt or Greenhouse-Geisser correction. Post hoc comparisons were performed using Holm’s method to identify specific frequency bands showing significant differences between stroke and control groups. We additionally compared the delta–beta power ratio to compare stroke patients and control subjects, using the same dilated lesion masks as in the previous analyses.

### Comparison among perilesional, disconnected and non-disconnected regions in stroke patients

We performed a repeated-measures ANOVA with location (perilesional, disconnected, and not-disconnected) and band (delta, theta, alpha, beta) as within-subject factors. Patients with lesions in brain regions that prevent the retrieval of cortical perilesional parcels (e.g., brainstem lesions), or that do not satisfy the disconnection criteria, were excluded from this analysis. Absolute EEG power was analyzed, as the repeated measure structure controls for stable inter-individual differences in overall signal amplitude.

Moreover, the spectral exponent (power-law slope) of the aperiodic component was estimated as a measure of changes in the frequency distribution of spectral power. Regional differences in the spectral exponent were evaluated using a repeated-measures ANOVA, with location (perilesional, disconnected, not-disconnected) specified as a within-subject factor.

### Relationship between EEG power in perilesional, disconnected and not-disconnected regions and behavioral measures in stroke patients

To investigate whether stroke severity was associated with relative power in different brain regions, we stratified patients into two groups for each frequency band. Low power (patients with values below the median) and High power (patients with values above the median). The grouping was performed based on the mean relative power computed within three ROI categories: perilesional, disconnected, and not-disconnected ROIs. We used the Wilcoxon rank-sum test to assess whether the 24h-NIHSS scores differed significantly between the two groups (FDR-corrected).

To investigate the relationship with behavioral measures from the neuropsychological battery, we derived four composite scores: executive functions (FAB, TMT-A, TMT-B), language (ENPA), memory (RAVLT, BVMT), motor (sum of NIHSS arm-left, leg-left, arm-right, and leg-right). Test-specific variation in sample size was linked with different ratio of completion of the cognitive tests. Each subtest was z-scored (relative to the controls). For NIHSS scores-scoring was performed within the patient sample, as NIHSS was not collected in controls, and reversed for congruence with the other cognitive tests (higher score indicates a better performance). First, we performed a multivariate analysis by means of PLS canonical correlation. Each power band considered only in the perilesional and disconnected regions contrasted with the four behavioral domains. Each mode was assessed for significance through a permutation test (n=1000). We considered only patients with all cognitive domain available for the PLS. Loadings from significant mode were plotted for interpretation. Second, we computed univariate Spearman correlations between each frequency band in perilesional and disconnected regions. Analyses for the PLS considered only patients with the whole cognitive domain spectrum available, while univariate analysis (NIHSS and behavioral scores) considered the maximum number of subjects available for each comparison. We used relative EEG power when correlating with behavioral measures to normalize for inter-individual differences in overall signal amplitude.

### Sensitivity Analyses

We performed a series of control analyses (Supplementary Table 8 to 13). First, we compared eyes-closed and eyes-open mean power values in all the areas using a repeated-measures ANOVA on absolute power across all regions, as relative power is confounded by condition-dependent physiological differences in alpha activity. Second, we repeated the repeated-measure Anova on absolute power across perilesional, disconnected and not-disconnected areas using an alternative probability threshold for the disconnection (40% and 60%). Third, the repeated-measure ANOVA assesses the effect of perilesional diameter (from 4 to 20 mm) on absolute power for each frequency band separately.

## DATA AVAILABILITY

All data reported in the present study are available from the authors upon reasonable request and all the software and algorithms used in the present study are cited in the Materials and methods section.

## ACKNOWLEDGEMENTS

The principal investigators wish to thank the participants who participated in the study for their time and effort.

## FUNDING

MC was supported by the Italian Ministry of Health “Eye-movement dynamics during free viewing as biomarker for assessment of visuospatial functions and for closed-loop rehabilitation in stroke” (EYEMOVINSTROKE; RF-2019-12369300); HORIZON-ERC-SyG (Grant No.101071900) “Neurological Mechanisms Of Injury And Sleep-Like Cellular Dynamics (NEMESIS)”; HORIZON-INFRA-2022 SERV (Grant No. 101147319) “EBRAINS 2.0: A Research Infrastructure to Advance Neuroscience and Brain Health”; LP and MC were funded from Cariparo Foundation Excellence grant 2023-2024 (Agreement No. 68076; AWARENET project); Ministry of University and Research, PRIN 20229Z7M8N “Tracking the brain priors for predictive coding”. LP is supported by the Associazione Italiana Ricerca Alzheimer Onlus (Airalzh-Grants-for-Young-Researchers - AGYR 2025). MVSV was supported by the INFRASLOW PID2023-152918OB-I00 financed by MICIU / AEI / 10.13039/501100011033/FEDER, UE.

## COMPETING INTERESTS

The authors report no conflicts of interest. All co-authors have seen and agree with the contents of the manuscript and there is no financial interest in reporting. We certify that the submission is original work and is not under review at any other publication.

## Notes

### Competing Interest Statement

The authors have declared no competing interest.

