## Supplementary Material for "White-matter disconnection shapes distributed cortical spectral dynamics after stroke"

### Supplementary Methods

**Neuropsychological Assessment**
Behavioral assessment was conducted at the time of EEG acquisition and consisted of a comprehensive battery of standardized neuropsychological tests. The assessment required approximately 90 minutes and was conducted in a quiet room. The complete neuropsychological battery included 11 different neuropsychological tests.

**Global cognitive screening.** The Oxford Cognitive Screen (OCS) provided a domain-specific profile of post-stroke cognition, designed to remain administrable in the presence of aphasia and neglect and covering attention and executive function, language, memory, number processing and praxis. The Frontal Assessment Battery (FAB) screened frontal-lobe function across conceptualization, mental flexibility, motor programming, sensitivity to interference, inhibitory control and environmental autonomy.

**Mood.** Depressive symptomatology was assessed with the Geriatric Depression Scale (GDS).

**Social cognition.** The Facial Expressive Action Stimulus Test (FEAST) assessed face memory, configural and part-based face processing, and recognition of facial expressions. The Story-based Empathy Task (SET) assessed mentalizing, dissociating the attribution of intentions and of emotions from non-social causal inference.

**Memory.** Verbal episodic memory was assessed with the Rey Auditory Verbal Learning Test (RAVLT), yielding measures of immediate span, learning across trials, delayed recall and recognition. The visuospatial analogue was provided by the Brief Visuospatial Memory Test (BVMT), with learning, delayed recall and recognition indices.

**Executive function and attention.** The Trail Making Test was administered in both forms, with part A indexing visual search and psychomotor speed and part B additionally loading on set-shifting. Covert orienting of visuospatial attention was measured with a Posner cueing paradigm, providing validity effects and an index of disengagement cost.

**Motor function.** Manual dexterity and fine motor speed were quantified with a pegboard task (PEG), administered to each hand separately.

**Language.** Language was evaluated with the Esame Neuropsicologico per l'Afasia (ENPA), an Italian battery assessing comprehension and production at the word and sentence level across oral and written modalities, including repetition, naming, reading and wri

Of the administered battery, two tests (i.e., PEG, Posner) were excluded due to insufficient completion rates. The social cognition and mood measures (FEAST, SET, GDS) were not considered further, as they fall outside the scope of the present study.

**Supplementary Table S1. Radiological features of the stroke sample**. Lesion laterality, small vessel disease markers and qualitative assessment of lesion location is shown.

| | | |
|---|---|---|
| **Markers of small vessel disease** | | |
| | Fazekas score, mean ± SD | 0.6 ± 0.6 |
| | Number of lacunes, mean ± SD | 0.6 ± 1 |
| **Lesion laterality, n (%)** | | |
| | Left hemisphere | 29 (59) |
| | Right hemisphere | 20 (41) |
| **Lesion location, n (%)** | | |
| | Subcortical | 16 (33) |
| | Cortical | 10 (20) |
| | Cortico-subcortical | 18 (37) |
| | Midbrain | 3 (6) |
| | Cerebellum | 2 (4) |

**Supplementary Table S2. NIHSS 24 h - Perilesional Low and High Delta**. Results of the multiple linear regression model examining the association between NIHSS score at 24 hours and the predictor variables perilesional delta group (i.e., Low vs High), acute treatment, lesion volume and age. The table reports regression coefficients (β), standard errors (SE), t-values, and p-values for each variable included in the final model (N=48).

| Term | Estimate | Std. Error | t value | p value |
|---|---|---|---|---|
| Intercept | 3.397 | 3.205 | 1.060 | 0.295 |
| Group Peri Delta (Low Delta) | -3.290 | 1.274 | -2.584 | *0.013* |
| Acute Treatment (None) | -3.556 | 1.756 | -2.025 | *0.049* |
| Acute Treatment (rt-PA) | -1.579 | 2.340 | -0.675 | 0.503 |
| Acute Treatment (rt-PA+EVT) | -0.773 | 2.049 | -0.378 | 0.708 |
| Lesion Volume | 0 | 0 | 1.024 | 0.312 |
| Age | 0.058 | 0.042 | 1.394 | 0.171 |

**Supplementary Table S3. NIHSS 24 h - Disconnected Low and High Delta.** Results of the multiple linear regression model examining the association between NIHSS score at 24 hours and the predictor variables disconnected delta group (i.e., Low vs High), acute treatment, lesion volume and age. The table reports regression coefficients (β), standard errors (SE), t-values, and p-values for each variable included in the final model (N=43).

| Term | Estimate | Std. Error | t value | p value |
|---|---|---|---|---|
| Intercept | 1.816 | 2.914 | 0.623 | 0.537 |
| Group Disc Delta (Low Delta) | -2.809 | 1.062 | -2.645 | *0.012* |
| Acute Treatment (None) | -3.496 | 1.470 | -2.684 | *0.011* |
| Acute Treatment (rt-PA) | -1.316 | 2.247 | -0.586 | 0.562 |
| Acute Treatment (rt-PA+EVT) | -0.635 | 1.702 | -0.374 | 0.711 |
| Lesion Volume | 0.00 | 0.00 | 1.857 | 0.072 |
| Age | 0.076 | 0.040 | 1.892 | 0.067 |

**Supplementary Table S4. NIHSS 24 h - Perilesional Low and High Alpha.** Results of the multiple linear regression model examining the association between NIHSS score at 24 hours and the predictor variables perilesional alpha group (i.e., Low vs High), acute treatment, lesion volume and age. The table reports regression coefficients (β), standard errors (SE), t-values, and p-values for each variable included in the final model (N=48).

| Term | Estimate | Std. Error | t value | p value |
|---|---|---|---|---|

| | | | | |
|---|---|---|---|---|
| Intercept | 1.570 | 3.218 | 0.488 | 0.628 |
| Group Peri Alpha (Low Alpha) | 2.425 | 1.196 | 2.027 | *0.049* |
| Acute Treatment (None) | -3.831 | 1.802 | -2.127 | *0.040* |
| Acute Treatment (rt-PA) | -3.138 | 2.292 | -1.369 | 0.178 |
| Acute Treatment (rt-PA+EVT) | -1.336 | 2.112 | -0.633 | 0.530 |
| Lesion Volume | 0.00 | 0.00 | 1.692 | 0.098 |
| Age | 0.044 | 0.044 | 1.017 | 0.315 |

**Supplementary Table S5. NIHSS 24 h - Perilesional Low and High Beta.** Results of the multiple linear regression model examining the association between NIHSS score at 24 hours and the predictor variables perilesional beta group (i.e., Low vs High), acute treatment, lesion volume and age. The table reports regression coefficients (β), standard errors (SE), t-values, and p-values for each variable included in the final model (N=48).

| Term | Estimate | Std. Error | t value | p value |
|---|---|---|---|---|
| Intercept | 1.223 | 3.197 | 0.383 | 0.704 |
| Group Peri Beta (Low Beta) | 2.810 | 1.279 | 2.197 | *0.034* |
| Acute Treatment (None) | -4.251 | 1.796 | -2.367 | *0.023* |
| Acute Treatment (rt-PA) | -3.328 | 2.270 | -1.466 | 0.150 |
| Acute Treatment (rtPA+EVT) | -1.228 | 2.098 | -0.248 | 0.806 |
| Lesion Volume | 0.00 | 0.00 | 0.630 | 0.532 |
| Age | 0.055 | 0.043 | 1.312 | 0.197 |

**Supplementary Table S6**. **NIHSS 24 h - Disconnected Low and High Beta.** Results of the multiple linear regression model examining the association between NIHSS score at 24 hours and the predictor variables other beta group (i.e., Low vs High), acute treatment, lesion volume and age. The table reports regression coefficients (β), standard errors (SE), t-values, and p-values for each variable included in the final model (N=48).

| Term | Estimate | Std. Error | t value | p value |
|---|---|---|---|---|
| Intercept | 0.312 | 2.937 | 0.106 | 0.916 |
| Group Disc Beta (Low Beta) | 2.765 | 1.017 | 2.720 | *0.010* |
| Acute Treatment (None) | -3.917 | 1.464 | -2.676 | *0.011* |
| Acute Treatment (rt-PA) | -3.009 | 2.117 | -1.422 | 0.164 |
| Acute Treatment (rt-PA+EVT) | -0.235 | 1.713 | -0.135 | 0.894 |
| Lesion Volume | 0 | 0 | 1.514 | 0.139 |
| Age | 0.058 | 0.040 | 1.463 | 0.152 |

**Supplementary Table S7. NIHSS 24 h – Not-disconnected Low and High Beta.** Results of the multiple linear regression model examining the association between NIHSS score at 24 hours and the predictor variables other beta group (i.e., Low vs High), acute treatment, lesion volume and age. The table reports regression coefficients (β), standard errors (SE), t-values, and p-values for each variable included in the final model (N=49).

| Term | Estimate | Std. Error | t value | p value |
|---|---|---|---|---|
| Intercept | 0.998 | 3.022 | 0.330 | 0.743 |
| Group Other Beta (Low Beta) | 3.488 | 1.106 | 3.155 | 0.003 |
| Acute Treatment (None) | -3.895 | 1.696 | -2.297 | 0.027 |
| Acute Treatment (rt-PA) | -3.630 | 2.158 | -1.683 | 0.100 |
| Acute Treatment (rt-PA+EVT) | -0.539 | 1.990 | -0.271 | 0.788 |
| Lesion Volume | 0.00 | 0.00 | 0.954 | 0.345 |
| Age | 0.051 | 0.040 | 1.287 | 0.205 |

**Supplementary Table S8. Results of the repeated measures ANOVA examining the effect of Condition (i.e. OC vs OA) and Band on Frequency Power.** Post-hoc comparisons were performed with Holm's method. The table reports F-values, degrees of freedom (df), p-values, and effect sizes for each main effect and interaction. Significant differences identified in post-hoc tests are indicated.

| Cases | Sphericity Correction | Sum of Squares | df | Mean Square | F | p |
|---|---|---|---|---|---|---|

| | | | | | | |
|---|---|---|---|---|---|---|
| Condition | None | 0.238 | 1 | 0.238 | 4.959 | 0.030 |
| Band | None | 96.462 | 3 | 32.154 | 282.769 | < .001 |
| Condition * Band | None | 0.458 | 3 | 0.153 | 15.190 | < .001 |

***Post Hoc Comparisons – Condition * Band – Conditional on Band (OC vs OA)***

| Band | Mean Difference | SE | df | t | p |
|---|---|---|---|---|---|
| Delta | -0.014 | 0.037 | 50 | -0.386 | 0.701 |
| Theta | 0.070 | 0.024 | 50 | 2.879 | 0.006 |
| Alpha | 0.148 | 0.024 | 50 | 6.256 | <.001 |
| Beta | -0.011 | 0.023 | 50 | -0.481 | 0.632 |

**Supplementary Table S9.** Results of the control repeated-measures ANOVA on absolute power across frequency bands for 40% disconnection probability threshold. The analysis examined the effect of Band on absolute power across perilesional, disconnected, and not-disconnected regions.

| Cases | Sphericity Correction | Sum of Squares | df | Mean Square | F | p |
|---|---|---|---|---|---|---|
| Band | Greenhouse-Geisser | 116.553 | 2.3 | 49.808 | 200.57 | < .001 |
| Area | Greenhouse-Geisser | 0.960 | 1.8 | 0.541 | 5.802 | 0.006 |
| Band * Area | Greenhouse-Geisser | 0.184 | 2.8 | 0.065 | 2.812 | 0.045 |

***Post Hoc Comparisons – Band × Area – Conditional on Band***

| Band | Comparison | | Mean Difference | SE | df | t | p |
|---|---|---|---|---|---|---|---|
| Delta | Perilesional | Disconnected | 0.022 | 0.039 | 43 | 0.564 | 0.576 |
| | | Not-disconnected | 0.133 | 0.049 | 43 | 2.712 | 0.019 |
| | Disconnected | Not-disconnected | 0.111 | 0.029 | 43 | 3.875 | 0.001 |
| Theta | Perilesional | Disconnected | 0.026 | 0.034 | 43 | 0.763 | 0.450 |
| | | Not-disconnected | 0.137 | 0.039 | 43 | 3.472 | 0.002 |
| | Disconnected | Not-disconnected | 0.111 | 0.029 | 43 | 3.882 | 0.001 |
| Alpha | Perilesional | Disconnected | 0.025 | 0.034 | 43 | 0.715 | 0.479 |
| | | Not-disconnected | 0.091 | 0.048 | 43 | 1.899 | 0.193 |
| | Disconnected | Not-disconnected | 0.067 | 0.039 | 43 | 1.730 | 0.193 |
| Beta | Perilesional | Disconnected | -0.015 | 0.025 | 43 | -0.588 | 0.860 |
| | | Not-disconnected | 0.026 | 0.033 | 43 | 0.797 | 0.860 |
| | Disconnected | Not-disconnected | 0.041 | 0.028 | 43 | 1.461 | 0.454 |

**Supplementary Table S10.** Results of the control repeated-measures ANOVA on absolute power across frequency bands for 60% disconnection probability threshold. The analysis examined the effect of Band on absolute power across perilesional, disconnected, and not-disconnected regions.

| Cases | Sphericity Correction | Sum of Squares | df | Mean Square | F | p |
|---|---|---|---|---|---|---|
| Band | Greenhouse-Geisser | 115.9 | 2.3 | 50.574 | 190.89 | < .001 |
| Area | Greenhouse-Geisser | 1.4 | 1.9 | 0.733 | 8.011 | < .001 |
| Band * Area | Greenhouse-Geisser | 0.2 | 3.1 | 0.067 | 2.921 | 0.036 |

***Post Hoc Comparisons – Band × Area – Conditional on Band***

| Band | Comparison | | Mean Difference | SE | df | t | p |
|---|---|---|---|---|---|---|---|
| Delta | Perilesional | Disconnected | -0.005 | 0.041 | 41 | -0.133 | 0.895 |
| | | Not-disconnected | 0.147 | 0.048 | 41 | 3.038 | 0.008 |
| | Disconnected | Not-disconnected | 0.153 | 0.033 | 41 | 4.601 | < .001 |
| Theta | Perilesional | Disconnected | 0.010 | 0.036 | 41 | 0.268 | 0.790 |
| | | Not-disconnected | 0.147 | 0.039 | 41 | 3.725 | 0.001 |
| | Disconnected | Not-disconnected | 0.137 | 0.033 | 41 | 4.107 | < .001 |
| Alpha | Perilesional | Disconnected | 0.006 | 0.039 | 41 | 0.151 | 0.881 |
| | | Not-disconnected | 0.108 | 0.048 | 41 | 2.260 | 0.058 |
| | Disconnected | Not-disconnected | 0.103 | 0.041 | 41 | 2.498 | 0.050 |
| Beta | Perilesional | Disconnected | -0.029 | 0.027 | 41 | -1.113 | 0.545 |
| | | Not-disconnected | 0.032 | 0.034 | 41 | 0.952 | 0.545 |
| | Disconnected | Not-disconnected | 0.062 | 0.031 | 41 | 1.983 | 0.162 |

**Supplementary Table S11** Results of the repeated-measures ANOVA assessing the effect of shell size (4–20 mm) on absolute power, conducted separately for each ROI (perilesional, disconnected, not-disconnected) on delta band (absolute power).

| Cases | Sphericity Correction | Sum of Squares | df | Mean Square | F | p |
|---|---|---|---|---|---|---|
| ROI | Greenhouse-Geisser | 2.215 | 1.6 | 1.382 | 7.572 | 0.002 |
| Shell Size | Greenhouse-Geisser | 0.001 | 1.2 | 0.001 | 1.018 | 0.335 |
| ROI * Shell Size | Greenhouse-Geisser | 0.011 | 2.1 | 0.006 | 3.093 | 0.049 |

***Post Hoc Comparisons – Shell size × ROI – Conditional on Shell size***

| Shell size | Comparison | | Mean difference | SE | df | T-score | p |
|---|---|---|---|---|---|---|---|
| 4-8 | Perilesion | Disconnected | -0.026 | 0.039 | 40 | -0.682 | 0.499 |
| | | Not- Disconnected | 0.132 | 0.052 | 40 | 2.555 | 0.029 |
| | Disconnected | Not- Disconnected | 0.158 | 0.033 | 40 | 4.768 | < .001 |
| 8-12 | Perilesion | Disconnected | -0.022 | 0.041 | 40 | -0.528 | 0.600 |
| | | Not- Disconnected | 0.132 | 0.052 | 40 | 2.539 | 0.030 |
| | Disconnected | Not- Disconnected | 0.153 | 0.033 | 40 | 4.685 | < .001 |
| 12-16 | Perilesion | Disconnected | -0.014 | 0.043 | 40 | -0.320 | 0.751 |
| | | Not- Disconnected | 0.133 | 0.052 | 40 | 2.568 | 0.028 |
| | Disconnected | Not- Disconnected | 0.147 | 0.034 | 40 | 4.348 | < .001 |
| 16-20 | Perilesion | Disconnected | 0.004 | 0.042 | 40 | 0.094 | 0.926 |
| | | Not- Disconnected | 0.142 | 0.050 | 40 | 2.859 | 0.013 |
| | Disconnected | Not- Disconnected | 0.138 | 0.034 | 40 | 4.078 | < .001 |

**Supplementary Table S12** Results of the repeated-measures ANOVA assessing the effect of shell size (4–20 mm) on absolute power, conducted separately for each location (perilesional, disconnected, other) on theta band (absolute power).

| Cases | Sphericity Correction | Sum of | df | Mean Square | F | p |
|---|---|---|---|---|---|---|
| ROI | Greenhouse-Geisser | 2.108 | 1.830 | 1.152 | 10.06 | < .001 |

| | | | | | | |
|---|---|---|---|---|---|---|
| Shell Size | Greenhouse-Geisser | 0.003 | 1.286 | 0.002 | 2.573 | 0.106 |
| Location * Shell Size | Greenhouse-Geisser | 0.004 | 1.729 | 0.002 | 1.013 | 0.359 |

**Supplementary Table S13** Results of the repeated-measures ANOVA assessing the effect of shell size (4–20 mm) on absolute power, conducted separately for each location (perilesional, disconnected, other) on alpha band (absolute power).

| **Cases** | **Sphericity Correction** | **Sum of** | **df** | **Mean Square** | **F** | **p** |
|---|---|---|---|---|---|---|
| ROI | Greenhouse-Geisser | 1.049 | 1.634 | 0.642 | 3.858 | 0.034 |
| Shell Size | Greenhouse-Geisser | 0.002 | 1.276 | 0.002 | 1.376 | 0.254 |
| Location * Shell Size | Greenhouse-Geisser | 0.001 | 1.700 | $8.744\times10^{-4}$ | 0.354 | 0.668 |

**Supplementary Table S14** Results of the repeated-measures ANOVA assessing the effect of shell size (4–20 mm) on absolute power, conducted separately for each location (perilesional, disconnected, other) on beta band (absolute power).

| **Cases** | **Sphericity Correction** | **Sum of** | **df** | **Mean Square** | **F** | **p** |
|---|---|---|---|---|---|---|
| ROI | Greenhouse-Geisser | 0.230 | 2.000 | 0.115 | 1.735 | 0.183 |
| Shell Size | Greenhouse-Geisser | 0.00 | 1.271 | $1.286\times10^{-5}$ | 0.020 | 0.932 |
| Location * Shell Size | Greenhouse-Geisser | 0.001 | 1.643 | $6.382\times10^{-4}$ | 0.450 | 0.601 |

**Supplementary Figure S1. Scatterplots showing the relationship between relative power and NIHSS stroke scale score across frequency bands and region categories**. Panels are organized by region (perilesional and disconnected) and frequency band (delta, theta, alpha, beta). In all panels, relative power is plotted on the x-axis and behavioral performance on the y-axis. Pearson correlation coefficients and corresponding uncorrected p-values are displayed within each plot.

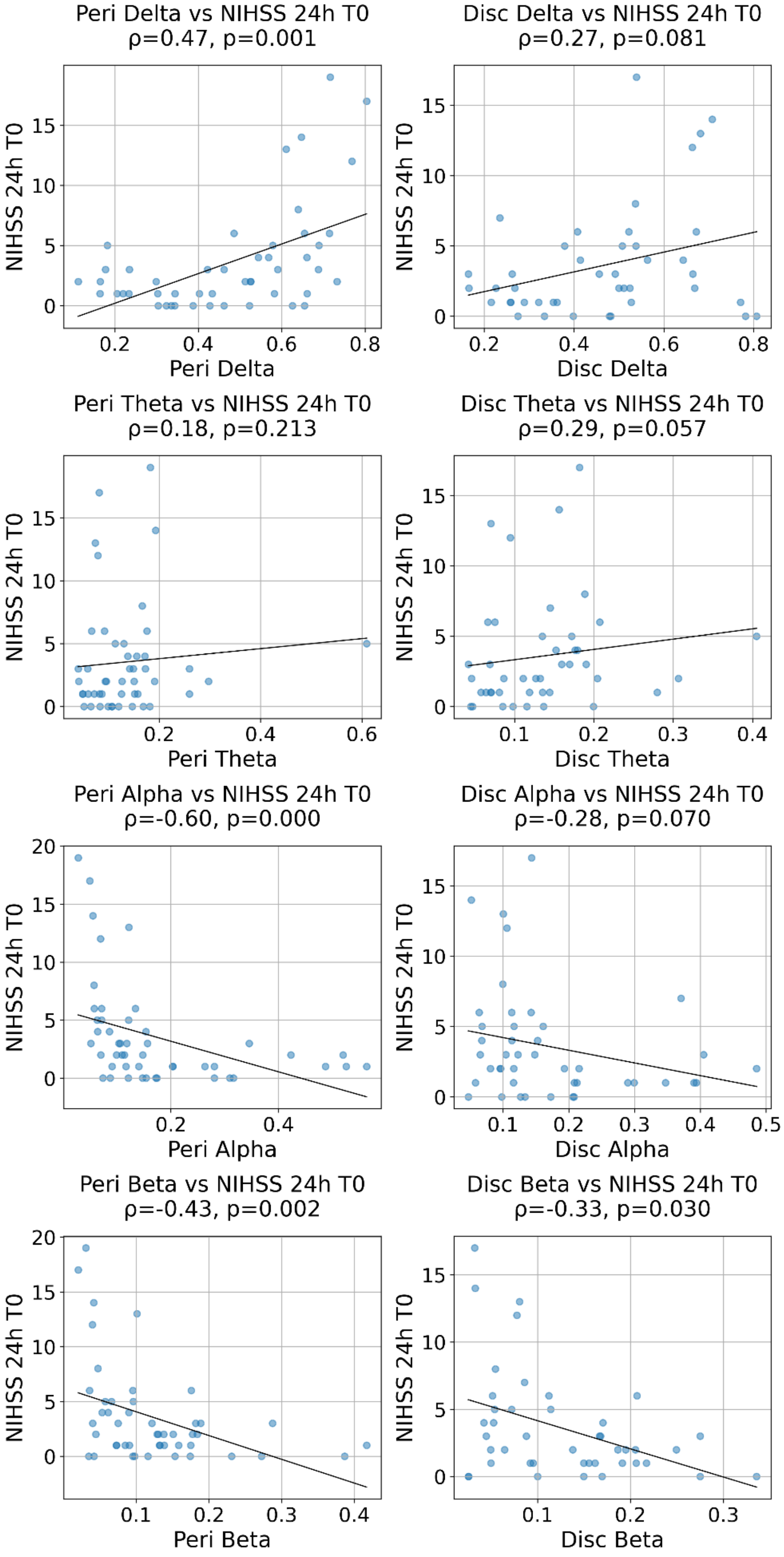

**Supplementary Figure S2. Scatterplots showing the relationship between relative power and executive functions composite score across frequency bands and region categories.** Panels are organized by region (perilesional and disconnected) and frequency band (delta, theta, alpha, beta). In all panels, relative power is plotted on the x-axis and behavioral performance on the y-axis. Pearson correlation coefficients and corresponding uncorrected p-values are displayed within each plot.

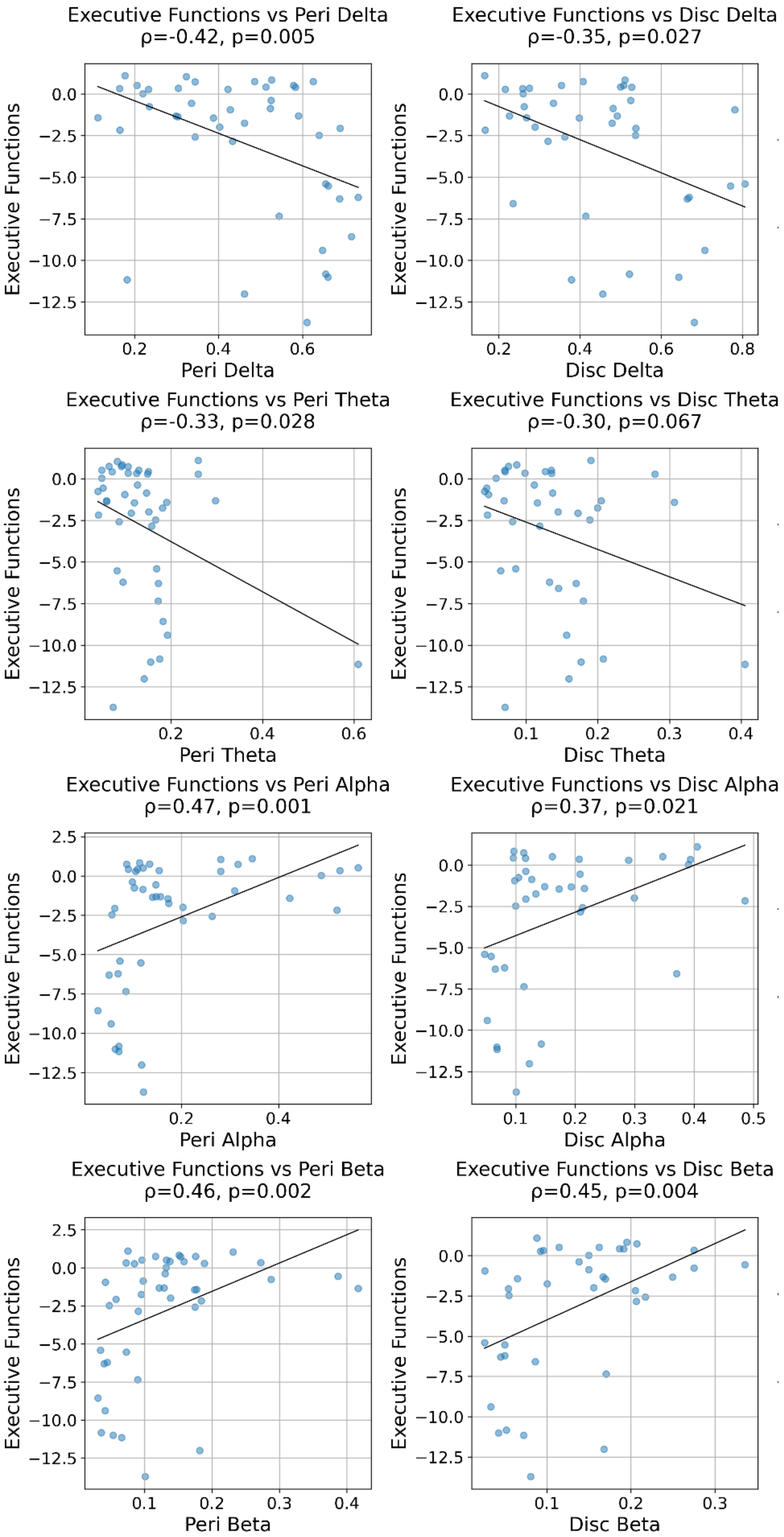

**Supplementary Figure S3. Scatterplots showing the relationship between relative power and language composite score across frequency bands and region categories.** Panels are organized by region (perilesional and disconnected) and frequency band (delta, theta, alpha, beta). In all panels, relative power is plotted on the x-axis and behavioral performance on the y-axis. Pearson correlation coefficients and corresponding uncorrected p-values are displayed within each plot.

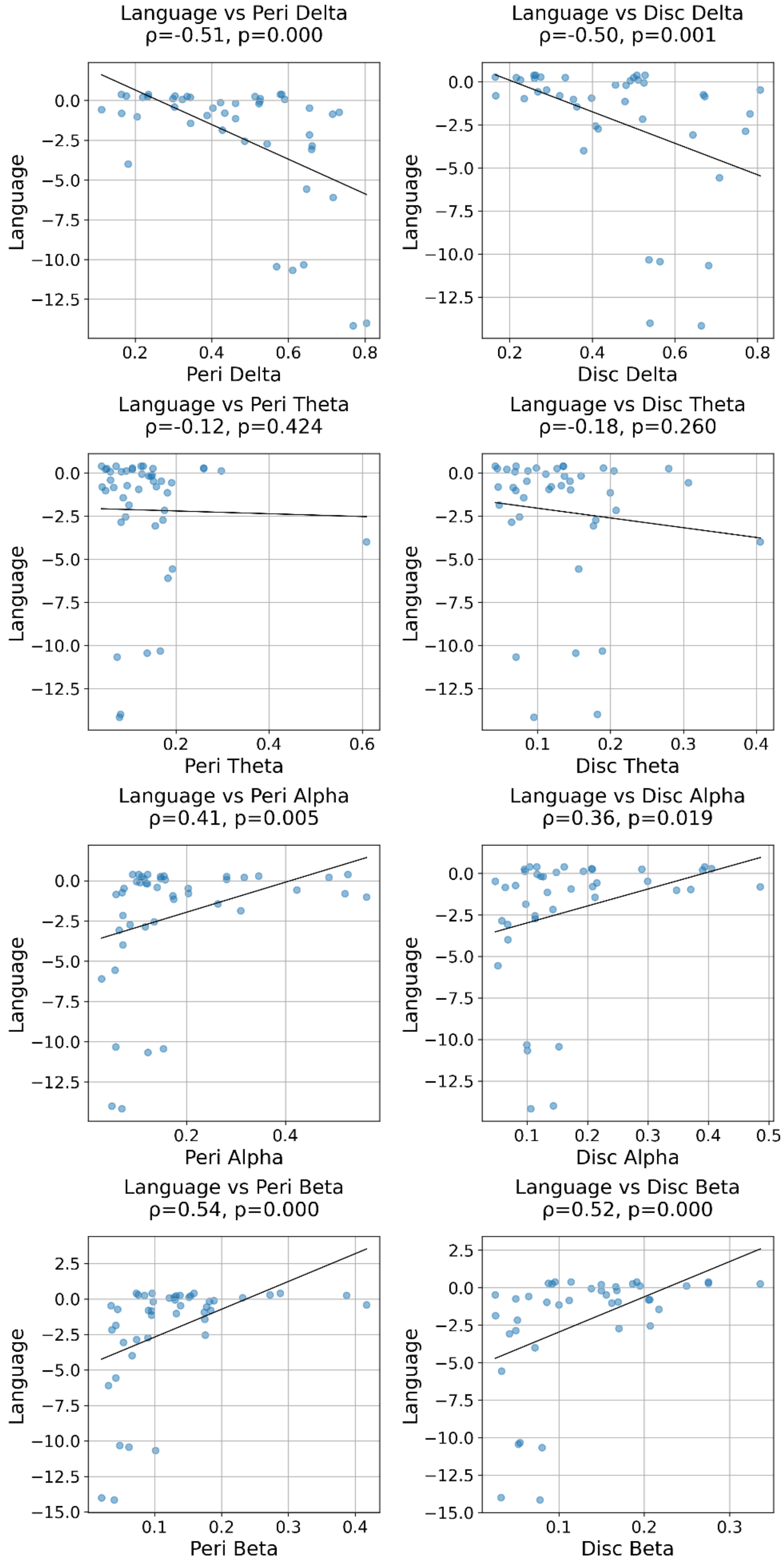

**Supplementary Figure S4. Scatterplots showing the relationship between relative power and memory composite score across frequency bands and region categories**. Panels are organized by region (perilesional and disconnected) and frequency band (delta, theta, alpha, beta). In all panels, relative power is plotted on the x-axis and behavioral performance on the y-axis. Pearson correlation coefficients and corresponding uncorrected p-values are displayed within each plot.

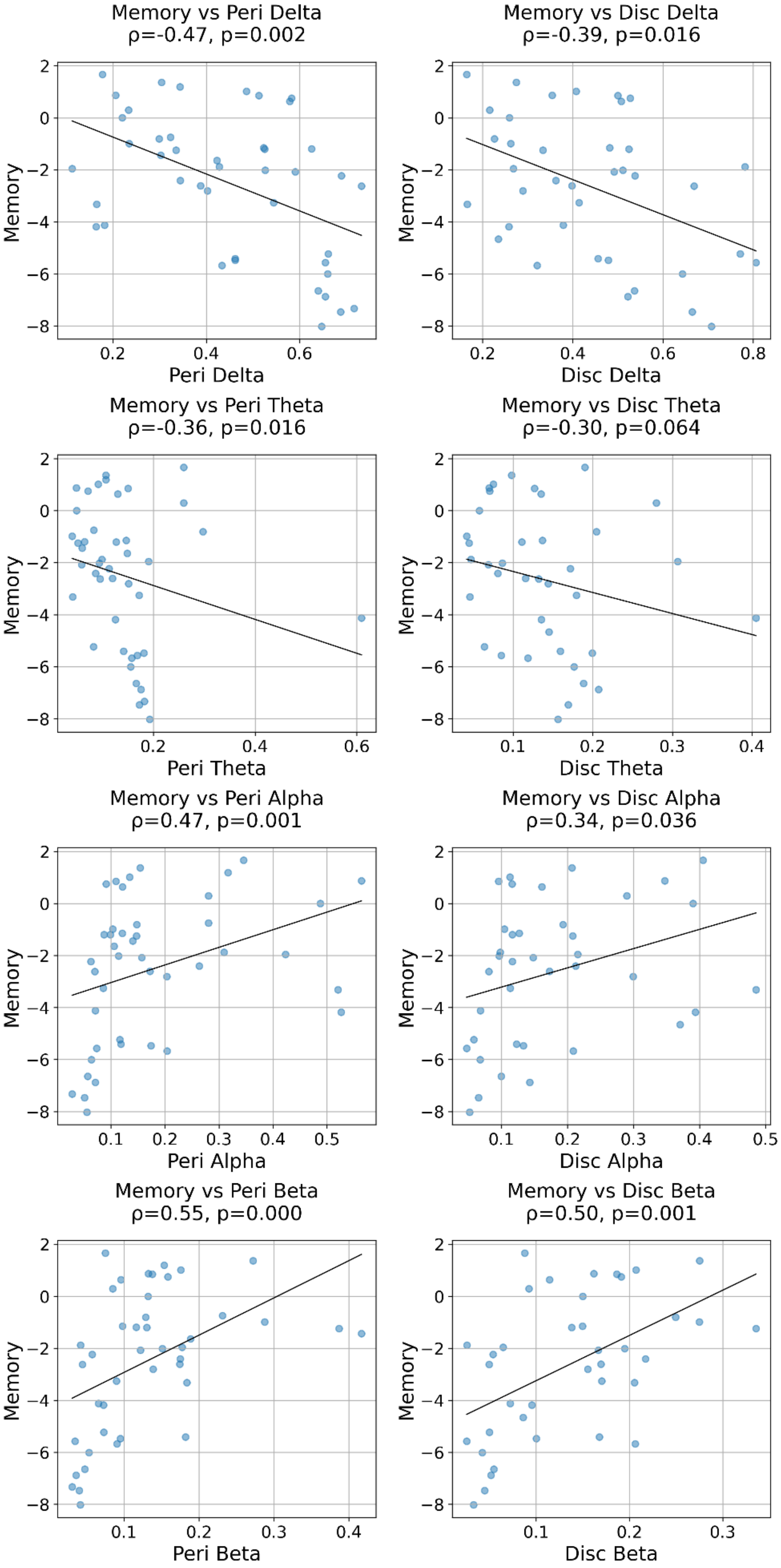

**Supplementary Figure S5. Scatterplots showing the relationship between relative power and motor composite score across frequency bands and region categories**. Panels are organized by region (perilesional and disconnected) and frequency band (delta, theta, alpha, beta). In all panels, relative power is plotted on the x-axis and behavioral performance on the y-axis. Pearson correlation coefficients and corresponding uncorrected p-values are displayed within each plot.

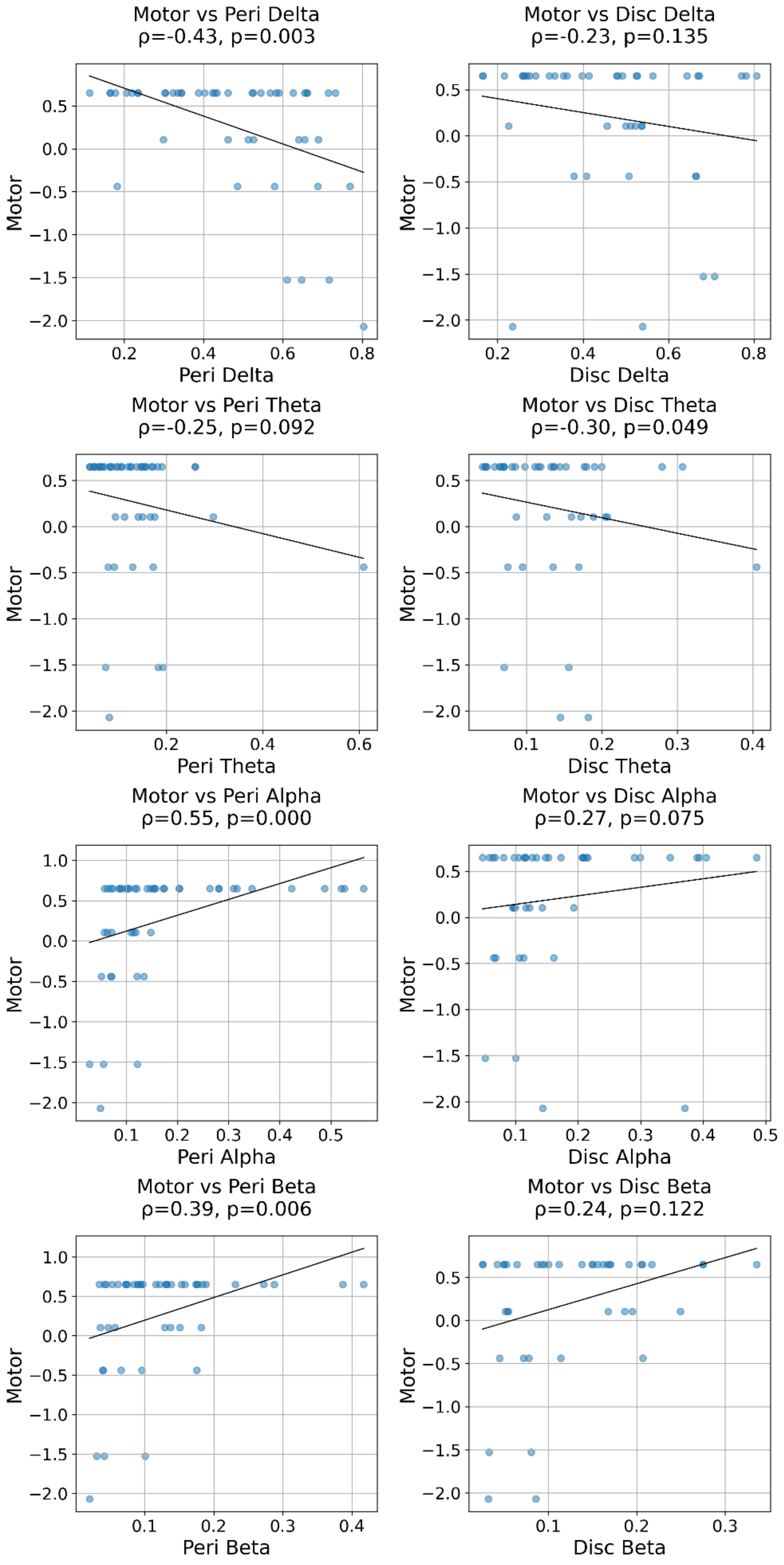

**Supplementary Figure S6.** Individual neuropsychological performance across tests. Each panel shows one test, dots are individual participants (jittered vertically for visibility), horizontal bars span the interquartile range and the filled circle marks the median. Scores are expressed as z-scores relative to the control group. Subscale scores were standardised individually before averaging into the composite shown. Subscales with zero variance in controls were excluded from the composites.

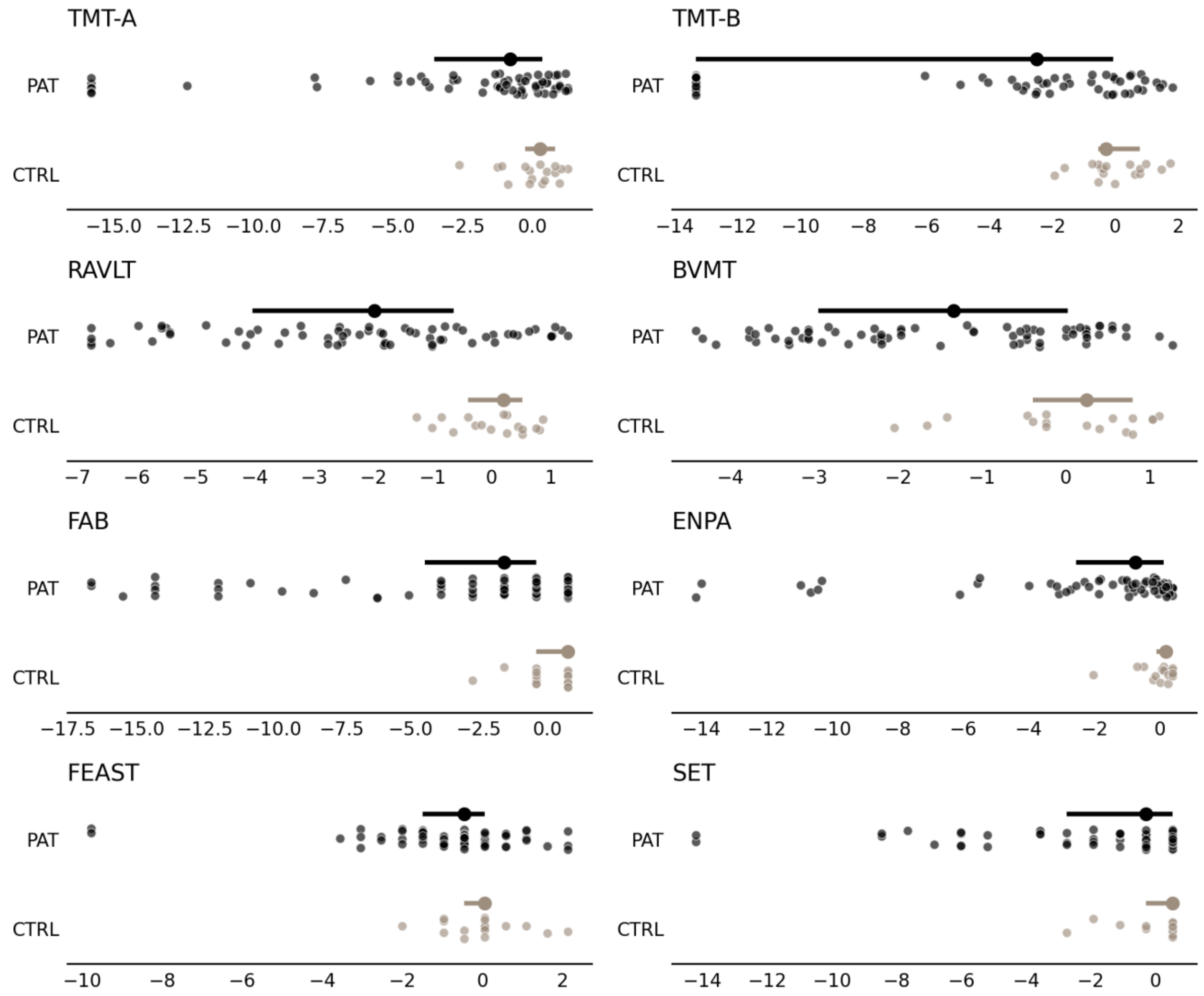
